# Fast, efficient and virus-free generation of *TRAC*-replaced CAR T cells

**DOI:** 10.1101/2021.02.14.431017

**Authors:** Jonas Kath, Weijie Du, Bernice Thommandru, Rolf Turk, Leila Amini, Maik Stein, Tatiana Zittel, Stefania Martini, Lennard Ostendorf, Andreas Wilhelm, Levent Akyüz, Armin Rehm, Uta E. Höpken, Axel Pruß, Annette Künkele, Ashley M. Jacobi, Hans-Dieter Volk, Michael Schmueck-Henneresse, Petra Reinke, Dimitrios L. Wagner

**Affiliations:** BIH Center for Regenerative Therapies (BCRT), Berlin Institute of Health (BIH), Berlin, Germany; Berlin Center for Advanced Therapies (BeCAT), Charité - Universitätsmedizin Berlin, corporate member of Freie Universität Berlin, Humboldt-Universität zu Berlin, and Berlin Institute of Health (BIH), Berlin, Germany; Integrated DNA Technologies, Inc., Coralville, IA, 52241, USA; Department of Nephrology and Intensive Care Medicine, Charité - Universitätsmedizin Berlin, corporate member of Freie Universität Berlin, Humboldt-Universität zu Berlin, and Berlin Institute of Health (BIH), Berlin, Germany; Deutsches Rheuma-Forschungszentrum (DRFZ), A Leibniz Institute, Berlin, Germany; CheckImmune GmbH, 13353 Berlin, Germany; Department of Translational Tumorimmunology, Max-Delbrück-Center for Molecular Medicine (MDC), 13125 Berlin, Germany; Department of Microenvironmental Regulation in Autoimmunity and Cancer, Max-Delbrück-Center for Molecular Medicine (MDC), 13125 Berlin, Germany; Institute of Transfusion Medicine, Charité - Universitätsmedizin Berlin, corporate member of Freie Universität Berlin, Humboldt-Universität zu Berlin, and Berlin Institute of Health (BIH), Berlin, Germany; Department of Pediatric Oncology and Hematology, Charité - Universitätsmedizin Berlin, corporate member of Freie Universität Berlin, Humboldt-Universität zu Berlin, and Berlin Institute of Health (BIH), Berlin, Germany; Institute of Medical Immunology, Campus Virchow-Klinikum, Charité - Universitätsmedizin Berlin, corporate member of Freie Universität Berlin, Humboldt-Universität zu Berlin, and Berlin Institute of Health (BIH), Berlin, Germany

## Abstract

Chimeric Antigen Receptor (CAR) redirected T cells are a potent treatment option for certain hematological malignancies. Recently, site-specific insertion of CARs into the T cell receptor (TCR) alpha constant (*TRAC*) locus using gene editing and adeno-associated viruses was shown to generate CAR T cells with improved functionality over their retrovirally transduced counterparts. However, the development of viruses for gene transfer is complex and associated with extensive costs at early clinical stages. Here, we provide an economical and virus-free method for efficient CAR insertion into the *TRAC* locus of primary human T cells *via* CRISPR-Cas mediated homology-directed repair (HDR). While the toxicity induced by transfected double-stranded template (donor) DNA was not fully prevented by pharmacological means, the combination of DNA-sensor inhibitors and HDR enhancers resulted in highly efficient gene editing with TCR-to-CAR replacement rates reaching up to 68%. The resulting TCR-deficient CAR T cells show antigen-specific cytotoxicity and cytokine production *in vitro*. Our GMP-compatible non-viral platform technology lays the foundation for clinical trials and fast-track generation of novel CAR T cells applicable for autologous or allogeneic off-the-shelf use.

## Introduction

Chimeric Antigen Receptor (CAR) redirected T cells are a potent treatment option for certain hematological malignancies, such as relapsed acute B-cell leukemia and aggressive B-cell lymphoma^1,2^. Clinical studies have shown CAR T cell efficacy in patients with treatment refractory multiple myeloma^3,4^. Furthermore, CAR T cell therapy is under development for solid tumors^5,6^. Their potential has been expanded to other disease entities and they have shown preclinical evidence in liver and cardiac fibrosis^7,8^, in autoimmune diseases ^9,10^ and in the prevention and treatment of allogeneic immune responses via CAR-reprogrammed regulatory T cells in transplantation medicine^11,12^.

Lenti- and retroviral gene transfer are the current gold standards for clinical manufacturing of CAR T cells. High expression levels after random transgene integration are achieved by the use of strong viral promoters. Excessive CAR expression, however, may contribute to tonic signaling and potentiate T cell susceptibility to activation induced cell death^13,14^ and exhaustion^15^. Furthermore, virus production is a laborious procedure requiring regulatory approval for preclinical research in many countries. Moreover, the costs associated with the production of clinical grade retroviruses are a limiting factor for the translation of new CAR T cell therapies^16^.

Preclinical evidence suggests that targeted integration of CAR transgenes into the T cell receptor (TCR) alpha constant chain (*TRAC*) gene locus allows for optimally regulated expression and superior functionality through reduced T cell exhaustion *in vitro* and *in vivo*^15,17^. Additionally, resulting *TRAC*-integrated and thus TCR/CD3-deficient CAR T cells avoid the risk of hazardous allogeneic graft-versus-host disease and offers a potential route towards off-the-shelf products after purification of TCR-negative CAR-T cells^18^. Furthermore, site-specific insertion of CAR transgenes using genome-editing techniques reduces the risk of insertional mutagenesis which can occur through retroviral gene transfer and it can avoid CAR expression in contaminating leukemia cells^19^.

*TRAC*-integration of CAR expression-modules has been achieved through nuclease-assisted homology directed repair (HDR)^15,17^. In previous studies, *Streptococcus pyogenes* (Sp)Cas9-single guide (sg)RNA ribonucleoprotein (RNP) complexes were transfected into anti-CD3/CD28-stimulated T cells followed by transduction with recombinant adeno-associated virus serotype 6 (rAAV6) for HDR donor template (HDRT) delivery^15,17^. Other nucleases have also been used in this context to induce DNA double stand breaks to initiate HDR^20,21^. While RNP-based gene editing is commonly used to modify T cells^22^ and retrovirally-transduced CAR T cells^23,24^, few research publications have reported the use of T cells with CARs integrated at the *TRAC* locus^15,17,20,21,25,26^. The aforementioned studies have used methods requiring high titers of rAAV6 for donor delivery, thus necessitating virological expertise and labor-intensive production even at preclinical scale. Therefore, we conclude that there is a need for methods that circumvent the practical hurdles of virus production.

Non-viral reprogramming is an attractive alternative to the conventional approaches which circumvents the need of rAAV6 for effective donor DNA delivery^15,27^. Roth *et al*. achieved efficient site-specific transgene insertions by co-electroporation of CRISPR-Cas RNPs and single or double stranded DNA (dsDNA) HDR templates^28^. Small fluorescent tags (∼700 base pairs [bp]) were efficiently integrated into different genetic loci in primary human T cells^28^. However, this approach is less efficient when delivering therapeutically relevant transgenes for adoptive T cell transfer with lengths exceeding 1500 bp^28^. Integration rates of tumor-specific TCRs range from 5% (∼2100 bp insert)^29^ to 15% (1545 bp insert)^28,30^. Even under optimized conditions, significant cell toxicity of up to 70-80% still occurs^31^. To date, virus-free gene editing has not been used to introduce CAR transgenes into the *TRAC* gene locus.

The toxicity of non-viral reprogramming in T cells is likely dependent on 2 major factors: The first is the physical strain of co-electroporation of dsDNA and RNP aggregates into cells. This issue has been partially addressed by the use of anionic polymers such as Poly-L-glutamic acid (PGA), which was recently shown to improve knock-in rates and reduce toxicity by physically dispersing large RNP aggregates into smaller complexes^32^. Additionally, Nguyen and colleagues suggested a modification of HDR-donor DNA using truncated Cas9 target sequences (tCTS) flanking both homology arms (HAs). The tCTS create binding sites for the co-delivered RNPs. The bound RNPs shuttle the DNA into the cell nucleus via the nuclear localization sequences (NLS) of the SpCas9 nuclease, leading to improved knock-in rates, however, it also increases toxicity when higher amounts of tCTS-modified donors are used^32^. The second major factor in the toxicity of non-viral reprogramming of T cells are the endogenous immune responses triggered by cytosolic dsDNA mediated by innate DNA-sensor protein pathways^33^ such as cyclic GMP-AMP synthase (cGAS), which is activated in a DNA length dependent manner^34^ promoting apoptosis *via* its downstream signaling effector stimulator of interferon genes (STING)^35^. To the best of our knowledge, interference with DNA-sensing has not been investigated to modulate gene editing outcomes in human T cells.

In this study, we provide a guide to fast and efficient virus-free integration of CARs into the *TRAC* locus of primary human T cells using an optimized method adapted from prior works^28,32^. We hypothesized that cell viability is primarily compromised by toxicity of dsDNA donor templates. We therefore assessed if temporary inhibition of cytosolic DNA-sensors increases the viability and CAR insertion rates after DNA transfection. Additionally, we investigated whether HDR-enhancing substances could amplify HDR integration rates with low amounts of dsDNA donors and different nucleases. Finally, we show that transient dsDNA-sensor inhibition synergizes with two commercially available HDR enhancers for effective CAR T cell generation. *In vitro* assays provide proof-of-principle for the functionality of CD19- and BCMA-specific CAR T cells generated under optimized conditions for *TRAC* integration.

## Results

### Nuclease selection to insert a CD19-specific CAR via CRISPR-Cas9 assisted HDR

In order to perform optimization experiments with a clinically relevant transgene, we rationally designed a second generation CD19-specific CAR with an intermediate length IgG1 hinge for detection purposes, a CD28 costimulatory domain followed by the CD3 zeta effector chain (**Fig. 1 a, Suppl. table 1**). We chose the sgRNA targeting *TRAC* exon 1 with the fewest *in silico* predicted off-targets for our experiments^21^, since careful selection of gRNAs is required to avoid off-targets and to ensure a high level of safety for future clinical translation^36^. We modified the 5’ end of the sgRNA spacer sequence to further decrease the risk for off-target editing (here: 5’- G**<u>G</u>**GAATCAAAATCGGTGAAT-3’ instead of 5’- GAGAATCAAAATCGGTGAAT-3’). Empirical nomination of off-target activity using GUIDE-seq indicated 5 potential off-target sites for the original sgRNA, and identified only 1 off-target event for our target sequence modified sgRNA **(Suppl. Fig. 1)**. To insert a 2015 bp-sized CAR transgene in frame with *TRAC* exon 1, we created a donor template for HDR (**Fig. 1a**, DNA- and amino acid sequences disclosed in **Suppl. table 1**). The insert was flanked with symmetric homology arms (HA) (400 bp each) consisting of a P2A-self-cleavage site, the CD19-CAR cDNA and a bovine growth hormone (bGH) derived polyadenylation sequence (pA). Two types of HDRT were prepared as previously described: one with regular HAs (reg. HA) and one with tCTS-modified HAs (tCTS-HA)^32^ (**Suppl. table 1**).

**Figure 1:**
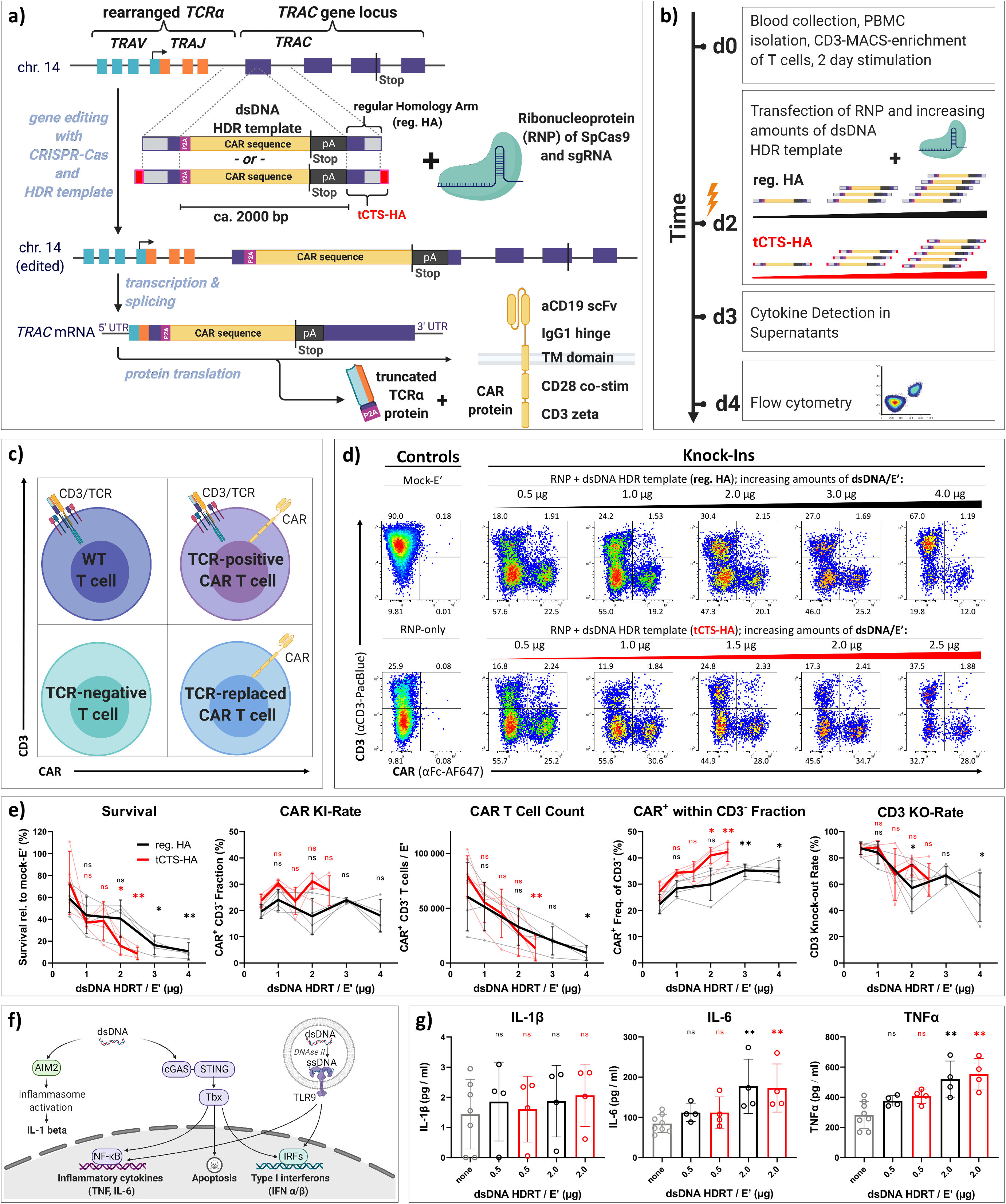
Low dosages of transfected dsDNA donor templates reduce toxicity during non-viral CAR T reprogramming. (a) Design of dsDNA donor templates for insertion of CAR transgenes into the *TRAC* locus. The transgene is comprised of a P2A self-cleavage site, the CAR encoding sequence followed by a STOP codon, and a bGH-derived poly-adenylation site (pA). Two template formats are used: the transgene flanked by regular homology arms (reg. HA, black) or homology arms with additional truncated Cas9 target sequences (tCTS-HA, red). (b) Experimental setup to evaluate co-transfection of RNPs and escalating doses of dsDNA donor templates with modified (tCTS-HA) and regular HA. (c, d) Representative flow cytometry plots show editing outcomes two days after co-transfection of RNP and escalating amounts of dsDNA donor templates of both HA formats. (e) Summary of flow cytometric analysis two days after transfection, n = 4 (two independent experiments). Black and red indicate the use of dsDNA HDRTs of regular HA or tCTS-HA format, respectively. Thick lines indicate mean values. Error bars indicate standard deviation. Lighter lines connect single data points from the same healthy donor over the different dsDNA amounts. Statistical analysis was performed using multiple Kruskal-Wallis tests with subsequent Dunn’s correction for multiple comparisons comparing values for each HDRT amounts with 0.5 µg HDRT as reference. (f) Illustration of common DNA-sensing pathways of the innate immune system as well as cytokines produced downstream. (g) Summary of supernatant analysis 24 hours after transfection of dsDNA HDRT for IL-1β (lower limit of detection (LOD): 0.85 pg/ml), IL-6 (lower LOD: 0.13 pg/ml) and TNFα (lower LOD: 0.05 pg/ml), n=4 (two independent experiments) Statistical analysis was performed using multiple Kruskal-Wallis tests with subsequent Dunn’s correction for multiple comparisons comparing cytokine concentrations for each HDRT amount with cytokine concentrations for electroporation without DNA. Asterisks represent different p-values calculated in the respective statistical tests (ns : p > 0.5; * : p < 0.05; ** : p < 0.01; *** : p < 0.001).

### Optimization of transfection conditions and pre-electroporation culture of T cells

First, we re-evaluated multiple parameters of the original protocols^28,32^ to generate an optimized version with increased cost-efficacy by substituting or reducing expensive reagents. In line with previous reports, 1×10^6^ activated primary human T cells per 20 µl electroporation solution were optimal for cell survival and knock-in rates (**Suppl. Fig. 2 a**). We found that the amount of Cas9 protein could be reduced from 10 to 4 µg / Electroporation (E’) without decreasing knock-in or knock-out efficiencies as determined by flow cytometry (**Suppl. Fig. 2 b**). Furthermore, we confirmed the positive effects of PGA on cell survival and relative CAR insertion rates for both dsDNA donor formats (regular HA and tCTS-HA) (**Suppl. Fig. 2 c**). When HDR donor templates were transfected without RNPs, no transgene expression was detectable by flow cytometry two days after electroporation (**Suppl. Fig. 2 d**). T cell stimulation with plate-bound anti-CD3/CD28 antibodies resulted in 29% (range: 27-41%) higher CAR^+^ T cell yield compared to anti-CD3/CD28-coated T cell activator beads^28,32^, without affecting insertion rates determined at day 7 post electroporation (**Suppl. Fig. 2 e**). The commercial P3 primary cell nucleofection buffer could be substituted with a self-made electroporation solution “1M”^37^ albeit leading to 2-5 %-points lower knock-in rates (**Suppl. Fig. 2 e**).

### Cell toxicity is dependent on the amount of transfected dsDNA donor templates

Using our optimized protocol (T cell stimulation in anti-CD3/CD28-coated wells, cell count: 1× 10^6^ / E’, amount of Cas9: 4 µg / E’, volume PGA: 0.5 µl / E’, buffer 1M), we investigated the effects of dsDNA HDR donor dosage and format as well as different pharmacological interventions on gene insertion rates and cell survival two days after transfection. In contrast to previous results^32^, we detected significant dose-dependent toxicity with regular HA dsDNA donors (**Fig. 1 b-e**). As expected, this trend was more pronounced with tCTS donor templates likely owing to enhanced DNA delivery as observed previously^32^. Mean toxicity at the highest HDRT amount tested (regular HA: 4 µg, tCTS: 2.5 µg) was 87% for regular HA and 90% for tCTS-HA, compared to mock-electroporated controls (**Fig. 1 d, e**). Generally, we observed moderate donor variability regarding DNA toxicity (**Fig. 1 e**). Within 48 hours, an average of 20% and 24% of living cells showed CAR surface expression after delivery of 0.5 µg HDRTs with regular HA and tCTS-HA, respectively (**Fig. 1 d, e**). Application of tCTS HDRTs tended towards increased CAR T cell counts at the lowest HDRT amount. In the resulting population of CD3-negative T cells, we observed an increase in relative CAR knock-in rates, up to 35% for regular HA and 42% with tCTS-HA at the respective highest amount of DNA used. However, overall CD3 knock-out rates decreased when using large HDRT amounts, indicating inhibitory DNA-RNP interaction with both HDRT-formats (**Fig. 1 e**). Cytokine analysis of cell culture supernatants collected 24 hours after transfection revealed dose-dependent release of IL-6 and TNFα, two cytokines that have been associated with intracellular sensing of dsDNA^33^ (**Fig. 1 f, g**). In contrast, IL-1β was barely above the minimal detection level and without a clear trend, indicating minimal AIM2-mediated inflammasome activation in T cells (**Fig. 1 f, g**). Taken together, dsDNA HDRT amounts exceeding 0.5 µg decreased CAR T cell yield through increased toxicity and reduced knock-out efficiency despite slightly increasing CAR knock-in rates at higher HDRT concentrations.

### Different DNA-sensor inhibitors can impact cell viability or CAR-integration rates

Transfected dsDNA activates innate immune pathways within T cells, thereby contributing to a loss in viability after electroporation of large dsDNA HDRT^34,35^ (**Fig. 1**). Aiming to improve T cell viability, we investigated whether temporary inhibition of intracellular dsDNA-sensors or associated downstream pathways could increase the survival of transfected T cells. Accordingly, we tested the effects of treating T cells before and/or after electroporation with one of the following compounds: 1. ODN A151, a modified DNA oligonucleotide, resembling the telomere repeat quadruplex 5’-TTAGGG-3’, which blocks endosomal Toll-like receptor 9 (TLR9) as well as other cytosolic DNA-sensors (cGAS, AIM2); 2. Ru.521, a small molecule antagonist of cGAS; 3. H151, a small molecule inhibiting STING; and 4. BX795, a small molecule that prevents activation of Tbx1, a kinase downstream of STING (**Fig. 2 a, b**). Cell viability after transfection of 2.0 µg of tCTS-modified HDRT was partially rescued by 6 hours pre-treatment with ODN A151 and BX795 (**Fig. 2 c**), but this effect was not observed with 1.0 µg reg. HA HDRT (**Fig. 2 d**). Extended presence of BX795 and RU.521, especially after electroporation, lead to minimal T cell recovery, indicating toxicity of continued exposure to these compounds. Strikingly, pre-treatment with the cGAS-inhibitor RU.521 appeared to increase the relative CAR insertion rate (**Fig. 2 c, d**).

**Figure 2:**
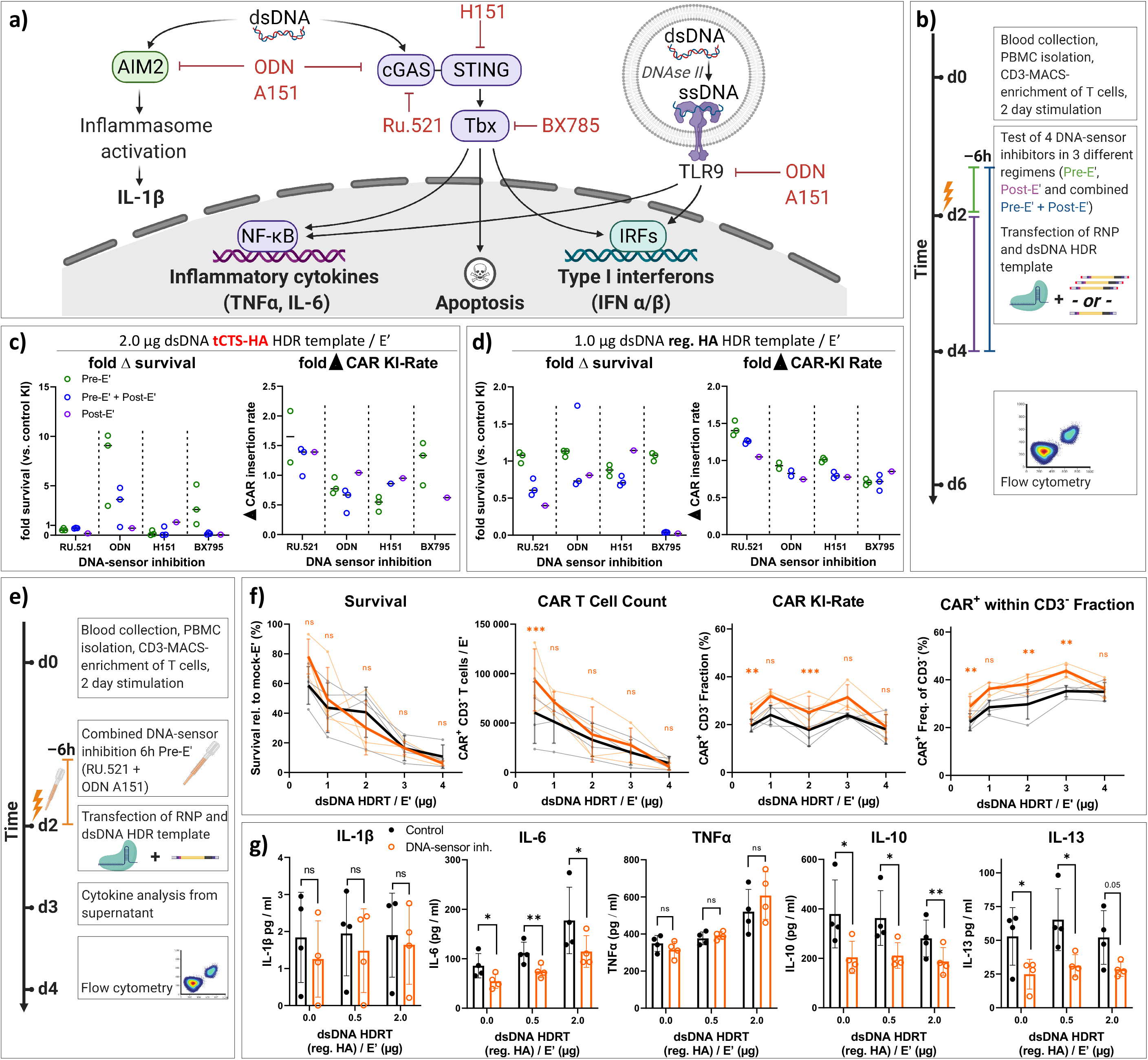
DNA-sensor inhibition increases relative CAR insertion rate with minimal improvement of survival at optimized dsDNA donor dosage. (a) Illustration of common DNA-sensing pathways in immune cells that induce downstream cytokine production, as well as the presumed mode-of-action of different DNA sensor inhibitors used in subsequent experiments. (b) Experimental setup to evaluate transient DNA-sensor inhibition using four different compounds. Green: Drug is added 6 hours before transfection. Purple: Drug is supplemented into the medium for 48 hours after transfection. Blue: Drug is added 6 hours before transfection and supplemented into the medium for 48 hours after transfection. (c, d) Relative survival and CAR-detection rate after co-transfection of T cells with RNP and high amount of tCTS-modified HDR template (c) or optimized dose of HDR template with regular HA (d) Both figures show the effects of differently timed interventions with each drug compared to editing results with non-treated control cells. These are set to 100% and indicated by dotted lines, (n=3). (e) Experimental set up for the combined addition of TLR-9 inhibitor ODN A151 and the cGAS-inhibitor RU.521 with escalating amounts of dsDNA donor templates (as in Fig. 1b). DNA sensor inhibitors were supplemented together into the medium 6 hours prior to co-transfection of RNP and reg. HA dsDNA donor templates. (f) Summary of flow cytometric analysis two days after transfection. Data were obtained in parallel to controls presented in Fig.1 d-g from four healthy donors in two independent experiments. Editing outcomes of T cells that received combined DNA-sensor inhibition prior to electroporation are shown in orange. Black indicates the control values from Fig. 1 e. Thick lines indicate mean values, error bars indicate standard deviation. Light dots represent individual values. Light lines connect these for each donor. Descriptive statistical analysis was performed using paired, two-tailed Student’s t tests comparing values for no intervention with values for DNA-Sensor inhibition. (g) Summary of supernatant analysis 24 hours after transfection for cytokines associated with DNA-sensing such as IL-1β (lower LOD: 0.85 pg/ml), IL-6 (lower LOD: 0.13 pg/ml) and TNFα (lower LOD: 0.05 pg/ml) as well as the Th2-associated cytokines IL-10 (lower LOD: 0.01 pg/ml) and IL-13 (lower LOD: 0.27 pg/ml) Descriptive statistical analysis was performed using paired, two-tailed Student’s t tests comparing cytokine concentrations for no intervention with cytokine concentrations for DNA-Sensor inhibition. Asterisks represent different p-values calculated in the respective statistical tests (ns : p > 0.5; * : p < 0.05; ** : p < 0.01; *** : p < 0.001).

### Treating T cells with two DNA-sensor inhibitors prior to electroporation improves CAR T cell generation at low dsDNA dose

Consequently, we evaluated whether a combined pre-treatment with ODN A151 and RU.521 could increase TCR-to-CAR replacement rates while retaining high T cell recovery with escalating amounts of reg. HA HDRT. (**Fig. 2 e, f**). We observed a small but consistent increase in the relative CAR expression rate as well as an increase in the absolute CAR T cell recovery at the lowest HDRT concentration (**Fig. 2 f**). Supernatant analysis revealed moderate suppression of IL-6 release by combined DNA-sensor inhibition, but no effects on TNFα and IL-1β, indicating an incomplete block in cytosolic dsDNA sensing. When transfection was performed with anti-CD3/CD28-stimulated peripheral blood mononuclear cells (PBMC), DNA-sensor inhibition reduced IL-1β production, which may be related to residual B cells and monocytes in the starting population (data not shown). Surprisingly, we saw a DNA-independent effect of DNA-sensor inhibition on Th2-associated cytokines IL-10 and IL-13 (**Fig. 2 g**), suggesting differential sensitivity of various T helper pathways. As cell cycle is a key determinant of DNA repair outcome^38^, we explored the effects of our drug combination on cell cycling prior to transfection as previously described^39^. Flow cytometric evaluation revealed an increased proportion of T cells within S-phase after 6 hours of pre-treatment with ODN A151 and RU.521 (**Suppl. Fig. 3**), which may explain the small increase in relative TCR-to-CAR replacement (**Fig. 2 f**). As DNA toxicity was not effectively prevented through transient DNA-sensor inhibition, we conclude that low dsDNA amounts (0.5 - max. 1.0 µg HDRT) are superior for effective CAR T cell generation (**Fig. 1e, 2f**).

### HDR-enhancers increase relative CAR-insertion rates in primary human T cells

Pharmacologic intervention in DNA repair processes represents an alternative strategy to obtain higher HDR rates as previously demonstrated for other gene editing procedures^40,41^. Therefore, we evaluated whether two commercially available HDR enhancing compounds (Alt-R HDR enhancer Version 1 [V1] and Alt-R HDR enhancer Version 2 [V2]) would increase the relative TCR-to-CAR replacement rate in our protocol (**Fig. 3 a**). We found a dose-dependent increase in the relative CAR insertion rate to 38% with V1 (range: 28-44%) and 43% with V2 (range: 31-53%) at their highest respective dose two days after electroporation (**Fig 3 b, c**). Due to the toxicity observed in some donors at the highest dose, we chose V1 at a concentration of 15 µM for subsequent experiments. The increase of CAR insertion mediated by use of HDR enhancer V1 was reproduced with another *TRAC*-HDR template^28^ and was increased irrespective of the chosen nuclease and sgRNA (**Fig. 3 d, e**). Suboptimal gene editing with an engineered Cas12a molecule from *Acidaminococcus species* (Alt-R A.s. Cas12a (Cpf1) Ultra, [Ultra-AsCpf1])^42^ as well as the HiFi SpCas9^43^ could be increased by the HDR enhancer V1 (**Fig. 3 d, e, f**).

**Figure 3:**
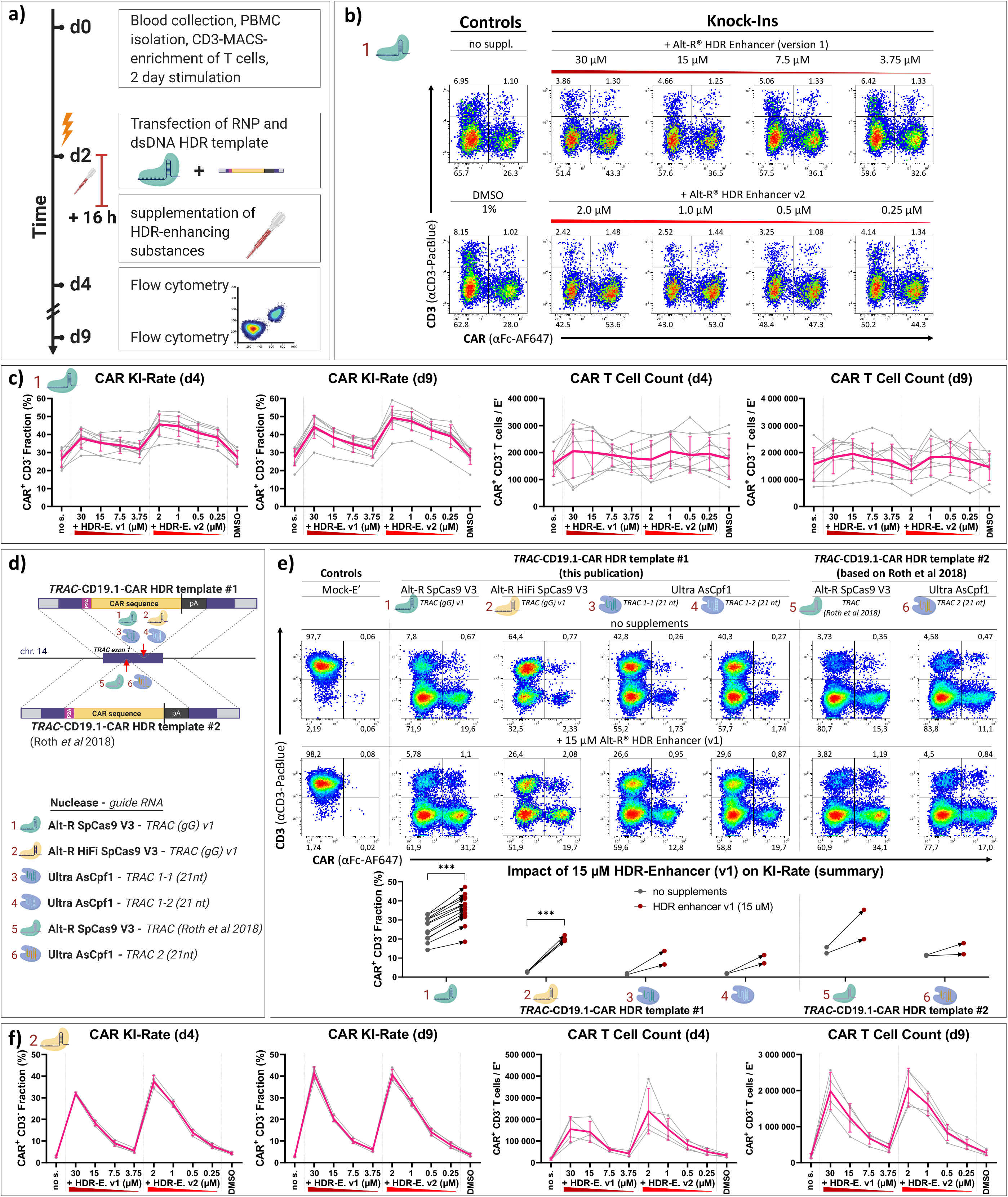
HDR enhancing compounds improve CAR integration rates in primary human T cells. (a) Experimental setup to identify the effects of HDR enhancing substances on *TRAC*-integration of CAR transgenes. (b) Representative flow cytometry plots comparing the effects of HDR enhancer V1 or V2, supplemented at different concentrations, on editing outcomes. (c) Summary of b with data from nine biological replicates in three independent experiments, two and seven days after transfection. The thick line indicates mean values. Light grey lines connect single data points, shown as light grey dots, from each biological replicate across the different conditions. Range of y-axes differs for CAR T cell count of d4 and d9. (d) Scheme illustrating two dsDNA donor templates of regular HA format for integration of CARs at two distinct sites of the *TRAC* locus, showing the three different Cas nuclease enzymes pre-complexed with different guide RNA to allow optimal targeting for each TRAC dsDNA donor template (=total of six distinct RNPs). (e) Representative flow cytometry plots and summary of results showing the effect of HDR enhancer V1 (15 µM) on CAR integration rates seven days after transfection with six different RNP / HDR template combinations as outlined in d. Statistical analysis was performed for RNP/HDRT combinations 1 and 2 using paired, two-tailed Student’s t tests comparing values for no intervention with values for HDR-Enhancer v1 (15 µM). (f). Test of Alt-R HiFi SpCas9 V3 Cas9 in an experimental setup analogous to a-c with a graphical representation as in c. A sketch of the RNP relates to d. Again, range of y-axes differs for CAR T cell count of d4 and d9. Data were obtained simultaneously from four biological replicates in one experiment. Asterisks represent different p-values calculated in the respective statistical tests (ns : p > 0.5; * : p < 0.05; ** : p < 0.01; *** : p < 0.001).

### Timing and temperature affects HDR modulation during virus-free CAR insertion in T cells

As gene editing with RNPs is likely to happen immediately after transfection^44–46^, we evaluated timing and cell culture conditions and their effects on pharmacologic modulation of CAR integration with HDR enhancers. HDR enhancers showed reduced effects when the supplemented medium was not pre-warmed, indicating their temperature-dependent activity (**Suppl. Fig. 4 a**). Of note, extended time between electroporation and transfer into HDR enhancer containing medium decreased the desired effects (**Suppl. Fig. 4 b**). Exposing T cells to HDR enhancers longer than the first three hours after transfection did not improve CAR integration rates (**Suppl. Fig. 4 c**). Both HDR enhancers improved CAR insertion (**Fig. 3 f**) and TCR/CD3 knock-out rates with HiFi SpCas9 in a dose-dependent manner (**Suppl. Fig. 4 d, e**). These results indicate that HDR enhancers can potently increase HDR-mediated CAR integration into the *TRAC* gene in primary human T cells.

### Transient DNA-sensor inhibition synergizes with HDR enhancers for efficient generation of TRAC-replaced CAR T cells

We subsequently tested whether the combination of DNA-sensor inhibition (ODN A151, Ru.521) and HDR enhancers could increase the absolute CAR T cell yield at 2 and 7 days after transfection (**Fig. 4 a**). We observed a synergistic effect on the relative TCR-to-CAR replacement rate as well as an increase in absolute CAR T cell recovery (**Fig. 4 b**). The increase in relative CAR insertion rates was independent of the donor template HA format (**Fig. 4 b**). In general, higher knock-in rates and improved CAR T cell yields were achieved when CD3-enriched T cells, instead of unselected PBMC, were used as a starting population (**Suppl. Fig. 5 a**). Synergy between DNA-sensor inhibition and HDR enhancer V1 was also observed after transfection of larger HDRT amounts, although the CAR T cell yield remained smaller than with optimized low dsDNA concentrations (**Suppl. Fig. 5 b**). In a further experiment using 1.0 µg reg. HA HDRT, temporary dsDNA-sensor inhibition further increased CAR integration rates with both HDR enhancers at all tested dosages, enhancing the CAR T cell yield after continued expansion (d9), but not at an early time point (d4) (**Fig. 4 c**). Importantly, neither pre-treatment with DNA-sensor inhibitors nor their combination with HDR enhancers significantly changed the CD4:CD8 ratio nor the T cell memory phenotype in comparison to mock-transfected T cells after 7 days of expansion (**Suppl. Fig. 6 a, b)**.

**Figure 4:**
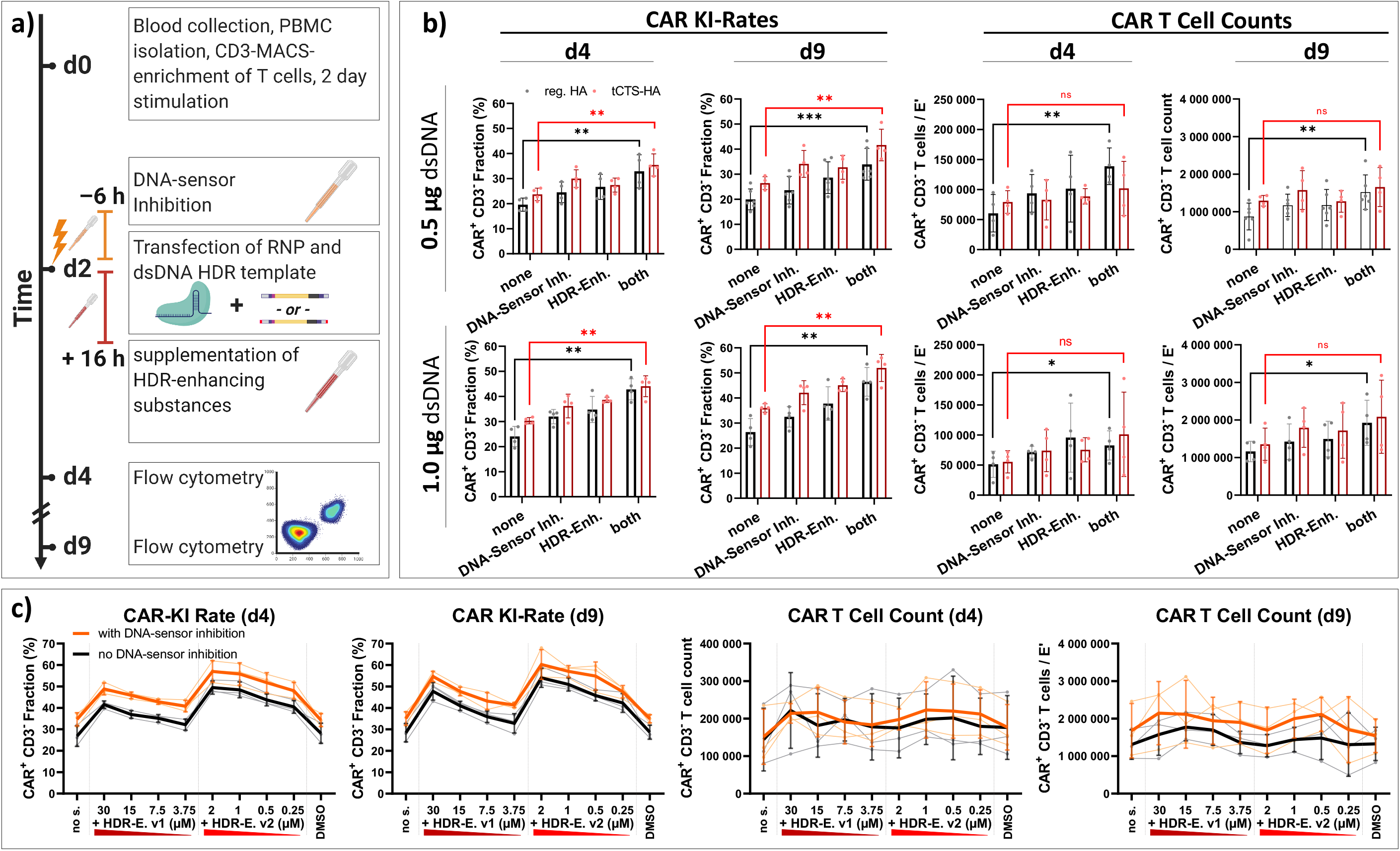
Synergy of DNA-sensor inhibition and HDR enhancers improves efficiency and yield of *TRAC-*replaced CAR T cell generation. (a) Experimental setup adding the combination of DNA-sensor inhibition and HDR-enhancement as an additional dimension to the setup originally presented in Fig. 1 e and Fig. 2 f. DNA-sensor inhibition was performed 6 hours prior to electroporation with the compounds ODN A151 and RU.521. After transfection, cells were cultured in HDR enhancer V1 supplemented medium (15 µM) for 16 hours. (b) Summary of relative CAR integration rates and CAR T cell counts two and seven days after transfection. 0.5 µg and 1.0 µg of dsDNA donor templates of both formats (reg. HA, black; and tCTS-HA, red) were used. Data were obtained in parallel to controls presented in Fig 1 d+e from four biological replicates. Furthermore, data from one additional experiment with two biological replicates only analyzed on d 9 were also included, mean +/-standard deviation. Statistical analysis was performed using paired, two-tailed student’s t test comparing values for no intervention (“none”) with values for a combined pharmacological intervention (“both”), individually for reg. HA and tCTS-HA. (c) Summary of CAR integration rates and CAR T cell yield comparing T cells with DNA-sensor inhibition (orange) or without it (black) prior to transfection of RNP with 1.0 µg reg. HA HDRT when cultured in medium supplemented with different dosages of the two HDR enhancers. Range of y-axes differs for CAR T cell count of d 4 and d 9. Data were obtained from three biological replicates in two independent experiments. Asterisks represent different p-values calculated in the respective statistical tests (ns : p > 0.5; * : p < 0.05; ** : p < 0.01; *** : p < 0.001).

### TRAC-replaced CAR T cells generated with virus-free method are functional in vitro and can be cryopreserved

Finally, we performed functional *in vitro* characterization of CAR T cells generated with our optimized protocol using the combined transient DNA-sensor inhibition with HDR enhancer V1. We designed HDR templates to introduce a CD19-specific CAR with a long extracellular spacer (CD19.2) or a B cell maturation antigen (BCMA)-specific CAR^47^. CD19.2-/BCMA-CAR T cells with CARs integrated into the *TRAC* locus were generated, expanded and cryopreserved at day 14. In line with previous reports^15^, CD19.2-CAR T cells with CARs integrated in the *TRAC* locus displayed a lower basal CAR expression in comparison to lentivirally transduced CD19.2-CAR T cells (**Suppl. Fig. 6)**. Similarly, site-specific insertion of a lentivirus-derived CD19.2-CAR overexpression cassette into the *AAVS1* safe-harbor locus induced higher CAR expression levels than *TRAC*-insertion (**Suppl. Fig. 6**). Prior to cryopreservation, antigen-specific cytotoxicity of *TRAC*-integrated CAR T cells was evaluated using the VITAL assay^48^. T cells were co-cultured with fluorescently labeled target antigen-expressing tumor cells (Nalm-6: CD19+, BCMA-; MM.1S: BCMA+, CD19-) and target antigen-negative control tumor cells (CD19-KO Nalm-6) (**Fig. 5 a**). We observed antigen-specific and dose-dependent lysis of the respective target cells within the 4-hour assay highlighting intact effector function (**Fig. 5 b, c**). Cryopreservation after CAR T cell manufacturing is a standard procedure to allow for quality control assessments, facilitate safe shipment after centralized production and to accommodate the individual patient treatment schedule. CAR T cells must retain effector function after a freeze-and-thaw cycle, especially in the case of allogeneic off-the-shelf approaches. Therefore, we evaluated the cytokine profile of thawed CAR T cells with TRAC integration of CARs after polyclonal stimulation with PMA/Ionomycin or tumor cell challenge (**Fig. 5 d, Suppl. Fig. 6 c, d**). As expected, PMA/Ionomycin stimulation induced a high percentage of cytokine producers among CD4^+^ and CD8^+^ wildtype T cells as well as the respective *TRAC*-integrated CD19.2- or BCMA-CAR T cell conditions (**Fig. 5 e-h**). In contrast, co-culture with CD19^+^ Nalm-6 tumor cells only induced cytokine production in CD19.2-CAR-positive CD4^+^ and CD8^+^ T cells (**Fig. 5 e-h**). Similarly, co-culture with BCMA^+^ MM.1S cells induced cytokine production selectively in BCMA-CAR expressing T cells (**Fig. 5 g, h**). Jurkat cells, which neither express CD19 nor BCMA, did not elicit cytokine production by CAR-expressing T cells (**Fig. 5 g, h**). Overall, these results indicate that CAR T cells with CARs integrated into the *TRAC* locus are functional after cryopreservation and thawing.

**Figure 5:**
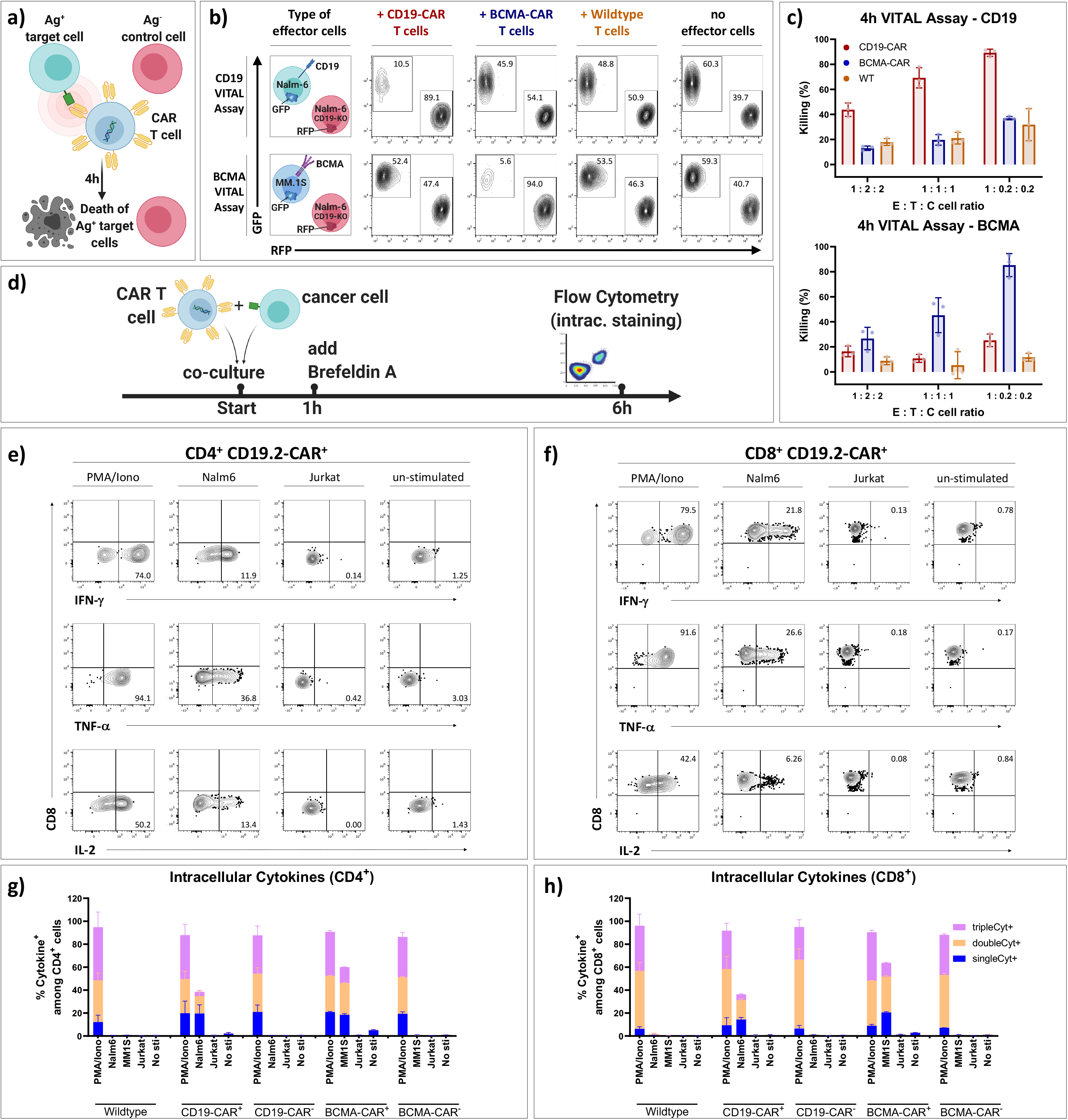
Functional characterization of *TRAC*-integrated CD19- and BCMA-specific CAR T cells *in vitro*. (a) Experimental setup of a flow cytometric VITAL assay. (b) Representative flow cytometry plots showing live target and control tumors cells after 4 hour VITAL cytotoxicity assays, shown for the highest effector (E):target (T):control (C) cell ratio tested (1 : 0.2 : 0.2). Either Nalm-6 cells (CD19^+^ BCMA^-^; top) or MM.1S cells (CD19^-^, BCMA^+^; bottom) were used as the target cell population. CD19 knock-out Nalm-6 cells served as controls in both cases. (c) Summary of VITAL assays performed simultaneously with edited cells from three healthy donors in three different E : T : C ratios, (n=3). (d) Experimental setup for detection of intracellular cytokines after different stimulations. (e,f) Representative flow cytometry plots of CD4^+^ (e) or CD8^+^ (f) CD19.2-CAR^+^ T cells after polyclonal stimulation (PMA/Ionomycin) or co-culture with either CD19^+^ Nalm-6 cells or CD19^-^ Jurkat cells. Representative gating strategy included in Suppl. Fig. 7. (g, h) Summary of e and f. Stimulations were performed in two technical replicates each. Data were obtained from three biological replicates of wildtype T cells, two biological replicates of *TRAC*-integrated CD19.2-CAR T cells and one biological replicate of *TRAC*-integrated BCMA-CAR T cells, each assessed in technical duplicates. Boolean gating was used to identify cells that produced one, two or three of the following cytokines: IFN-γ, IL-2 and TNFα. Error bars show standard deviation.

## Discussion

Here, we show that the efficient manufacturing of TCR-replaced CAR T cells is possible using a virus-free technique. The baseline insertion rates of a 2015 bp CAR transgene exceed those of previously published virus-free approaches with similarly sized inserts. We show that two different pharmacological interventions synergistically improve both the relative knock-in rate and the absolute CAR T cell yield. This method may also be applicable to improve orthotopic TCR replacement^29^, repurposing endogenous transcriptional pathways as proposed by others^25^ or even pooled knock-in screens, such as PoKi-seq^30^.

We have independently replicated findings by Nguyen *et al*. that the anionic nanoparticle PGA increases efficiency and decreases toxicity during electroporation of RNP and dsDNA HDRT, although the relative effects were smaller than originally reported^32^. Similarly, tCTS-modified dsDNA HDRTs displayed an advantage over the regular HA configuration regarding knock-in efficiency, although overall CAR-T cell yield was only improved when the tCTS-modified HDRT was transfected in small amounts (0.5 µg). These differences may result from other changes in our protocols, such as the significant reduction of the co-transfected RNP amount as well as the plate-bound stimulation^22^. The latter further reduces costs and may circumvent cell loss by bead depletion prior to transfection.

Increased amounts of transfected dsDNA donors resulted in substantially reduced cell survival, coincided with increased secretion of IL-6, and TNFα and decreased overall CAR T cell yields. Combined DNA-sensor inhibition (ODN A151, RU.521) partially reduced IL-6 release, but it did not affect TNFα secretion. In line with our initial hypotheses, these findings indicate that the toxicity of the virus-free approach is caused mainly by transfection of dsDNA donors, and is at least partially mediated *via* classical DNA sensing pathways. Independent of transfected DNA amount, our combined DNA-sensor inhibition reduced the release of Th2-associated cytokines such as IL-13 and IL-10, although the underlying mechanism remains to be elucidated. DNA-sensor inhibition provides a route towards further improvements of this method, potentially through inhibitors with higher potency or interference with additional (non-classical) DNA sensing pathways^49^.

Lowering the amount of transfected dsDNA donor may reduce toxicity by decreasing the physical strain imposed on the cell as well as provoking a weaker innate immune response. Whilst this is advantageous in terms of absolute CAR T cell yield, it also results in decreased relative HDR-rates. Thus, in order to increase CAR integration rates with low amounts of dsDNA donors, modulation of DNA repair towards HDR is desirable. Fortunately, multiple HDR enhancing drugs have been previously identified that are able to achieve this goal^40,41^. Our data show that two commercially available HDR enhancers increase relative knock-in rates in a dose-dependent manner. Intriguingly, suboptimal editing with a Cas12a derivative could be improved, which may be of interest for groups targeting AT-rich regions in the genome^50^. Suboptimal editing with HiFi SpCas9 is likely caused by the intended mismatch in our *TRAC* sgRNA and the low amount of RNP used, as others can achieve highly efficient editing with this nuclease and rAAV6^21,43^.

We discovered that pre-electroporation DNA-sensor inhibition treatment increased relative knock-in rates of the CAR constructs. As HDR is cell cycle dependent^38,51^, the improved knock-in rates seen here may in part be attributed to a small increase in the proportion of T cells within S-phase at the time of transfection. Alternatively, this may be directly related to the use of the cGAS-inhibitor, RU.521. Recent studies reported that nuclear cGAS is a general inhibitor of DNA repair and prevents homology-directed DNA repair through direct condensation of genomic DNA^52,53^. In particular, this may explain the synergy between cGAS-directed DNA sensor inhibition and HDR enhancers observed in our experiments. Before clinical application, HDR enhancers and DNA-sensor inhibitors must be carefully evaluated regarding their effects on off-target gene editing, genomic DNA integrity and homology-independent dsDNA donor integration events.

Along with the disclosed DNA sequences, the protocol is easily adaptable for preclinical investigations and after careful evaluation of safety, it may be scalable to suit clinical purposes in an allogeneic setting. It circumvents the need for the laborious, time consuming and costly manufacturing of viral vectors. By removing the practical hurdle of virus production, we hope our described technique may encourage groups to start working on CAR T cells, in particular *TRAC*-replaced CAR T cells. Our simplified technology platform could pave the way for the economical development of CAR T cell products and thus small scale CAR T cell trials. In doing so, we believe this platform may accelerate innovation for the treatment of various rare diseases as well as for solid tumour disease.

## Data availability statement

All underlying data of this manuscript can be obtained from the corresponding author upon request except the DNA sequence information of the BCMA-CAR (provided by A.R. and U.E.H.) and the CD19.2-CAR (from A.K.). Next-generation sequencing data from Guide-Seq experiment was deposited in the Sequencing Read Archive (SRA) repository of the National Center for Biotechnological Information (NCBI) under the Bioproject Accession Code : PRJNA701496.

## Contributions

J.K. designed this study, planned and performed experiments, analysed results, interpreted the data and wrote the manuscript. W.D. performed experiments, analysed results, interpreted data and edited the manuscript. B.T. and R.T. provided reagents, performed GUIDE-seq experiments, analysed results and interpreted data. L.Am., M.S., T.Z., S.M., L.O., A.W., L.Ak. performed experiments and analysed results. A.R. and U.E.H. provided reagents (BCMA-CAR, MM1S cell line), interpreted data and edited the manuscript. A.P. provided reagents and interpreted data. A.K. provided reagents (CD19.2-CAR, lentivirus, Nalm-6 cell line), interpreted data and edited the manuscript. A.J. planned and interpreted data and edited the manuscript. H.-D.V. provided reagents, interpreted data and edited the manuscript. M. S.-H. planned experiments, interpreted data and edited the manuscript. P.R. supervised the study, interpreted data and edited the manuscript. D.L.W. designed and led the study, planned and performed experiments, analysed results, interpreted data and wrote the manuscript. All authors discussed, commented and approved the manuscript in its final form.

## Acknowledgements

We would like to thank Dr. Alexander Marson and Dr. Theodore L. Roth for protocols on their original work on non-viral T cell reprogramming. We show gratitude to Lonza GmbH for temporarily providing the 4D-Nucleofector™ 96-well unit for the DNA-escalation experiment presented in Fig. 1e, 2e, 4b, Suppl. Fig.5, in particular we would like to thank Dr. Melanie Homberg and Dr. Nina Novak for technical assistance. We thank Jan Csupor for helping with multiplex-cytokine analysis of cell culture supernatants. The AAVS1 donor template was generously provided by Dr. Anne Floriane Hennig and Prof. Dr. Uwe Kornak (University of Göttingen). Norohiro Watanabe (Baylor College of Medicine, USA) provided insights for the cell cycle analysis protocols. We thank Dr. Ciaran M. Lee and Dr. Kathleen Anders for the critical revision of the manuscript. Finally, we would like to express our gratitude to Dr. Nicola R. Brindle for scientific language editing our manuscript.

## Methods

### GUIDE-Seq

GUIDE-Seq^54^ for unbiased identification of CRISPR-Cas off-targets was performed as previously described^43,55^. Briefly, SpCas9-expressing HEK293 cells were co-transfected with dsDNA oligonucleotides (GUIDE-seq tag, for sequence see^54^) and either the original^21^ or target-sequence modified *TRAC* sgRNA (this study). Subsequently, genomic DNA was isolated after 72 hours using QuickExtract (Lucigen, USA) and fragmented using the LOTUS™ DNA library kit (IDT, Coralville, USA). Libraries were generated according to the original protocol^54^, followed by Illumina based next generation sequencing. Read alignment and data analysis was performed based on GUIDE-seq software^54^.

### PBMC isolation and T cell enrichment

The study was performed in accordance with the declaration of Helsinki. Peripheral blood was obtained from healthy human adults after informed consent (Charité ethics committee approval EA4/091/19). Peripheral blood mononuclear cells (PBMC) were isolated using density-gradient centrifugation. Fresh heparinized whole blood was diluted 1:1 with sterile PBS (Gibco) and layered onto PANcoll separation medium (PAN Biotech) in 50 ml tubes (Falcon). Centrifugation was performed, with the brake function turned off, at 800 g for 20 mins. Subsequently, the mononuclear cell layer was harvested and diluted in sterile PBS. After an initial centrifugation at 450 g for 10 minutes, supernatant was discarded. The pellet was resuspended in sterile PBS and centrifuged at 300 g for 10 minutes. Afterwards, the supernatant was discarded. PBMC pellets were resuspended in 10-20 ml PBS and counted using a Neubauer hemocytometer. Unless otherwise stated, PBMCs were positively enriched for CD3^+^ T cells using magnetic column enrichment with human CD3 microbeads according to the manufacturer’s recommendations (LS columns, Miltenyi).

### Cell culture

PBMC or enriched T cells were cultured in RPMI 1640 (PAN Biotech), 10% heat-inactivated fetal calf serum (FCS) (Biochrom) and recombinant IL-2 (100 IU/ml), IL-7 (10 ng/ml) and IL-15 (5 ng/ml). T cell stimulation was performed for 48 hours on anti-CD3/28 coated tissue culture plates unless stated otherwise. Coating of vacuum gas plasma treated polystyrene 24-Well-Tissue-Culture plates (Corning Inc.) was performed overnight with 500 µl/well of sterile ddH2O supplemented with 1 µg/ml anti-CD3 mAb (clone OKT3, Invitrogen) and 1 µg/ml anti-CD28 mAb (clone CD28.2, Biolegend).Plates were washed twice in PBS and once in RPMI without letting the wells dry out. T cells were seeded at a density of 1-1.5× 10^6^ per 24-well. For some experiments, stimulation was performed with Dynabeads Human T-Activator CD3/CD28 beads (Invitrogen) according to the manufacturer’s protocol. Prior to nucleofection, beads were removed by incubation on strong magnet stands. The tumor cell lines Nalm-6 and MM.1S were obtained from the German Collection of Microorganisms and Cell Cultures GmbH (DSMZ) and cryopreserved for later use. The MM.1S cell line was genetically manipulated to overexpress GFP and firefly-luciferase (by A.R. and U.E.H.). All cell lines were freshly thawed and passaged 2-6 times prior to use in assays. Cell lines were cultured in RPMI1640 supplemented with 10% heat-inactivated FCS, 100 I.U./mL penicillin and 100 µg/ml streptomycin. Tumor cell lines were split every 2-3 days. All cell culture was performed at 37°C and 5% CO_2_.

### Generation of dsDNA donor template for homology-directed insertion of a Chimeric Antigen Receptor

Design of homology arms was performed as recently described^28^. Cloning of HDR donor templates with over 2000 bp insert size was performed with multiple fragment In-Fusion cloning according to the manufacturer’s protocol (Clontech, Takara). In brief, synthesis of 400 bp dsDNA sequences homologous to the targeted locus were commissioned (gBlocks, IDT) with a total overlap of 16 bp with either the insert or the pUC19 vector backbone. Similarly, inserts consisting of a P2A self-cleaving peptide, CAR/reporter transgene and a bovine growth hormone-derived polyadenylation sequence were designed and synthesis was commissioned (gBlocks, IDT). As a model insert for a therapeutically relevant transgene, a CD19-specific chimeric antigen receptor was designed based on the original FMC63 scFv. We chose an intermediate length IgG1 hinge (for staining purposes) and a CD28 transmembrane and costimulatory domain linked to a cytosolic CD3 zeta domain. The CAR sequence was rationally designed and subsequently codon-optimized using two separate algorithms: first, using the COOL-algorithm^56^; in a second step, IDT’s codon optimization tool was used to eliminate any complexities and allow DNA synthesis. In-Fusion cloning strategies were planned with SnapGene (from Insightful Science; snapgene.com). For other experiments, CAR transgenes were PCR amplified (Kapa Hotstart HiFi Polymerase Readymix, Roche) from lenti-/retroviral expression plasmids with primers that allowed In-Fusion reaction with the existing donor template backbones (*TRAC, AAVS1*). The CD19.2-CAR with a long IgG1 spacer, CD28 transmembrane, CD28 co-stimulatory domain and CD3 zeta was originally designed and cloned by A. Künkele. The BCMA.4-1BB-zeta CAR was previously described^47^, and the plasmid was provided by A.R. and U.E.H. (MDC, Berlin). In-Fusion reactions were performed in 5 µl reactions at the recommended volume ratios. 1 µl of In-Fusion reaction mixtures were transformed into Stellar Competent E. coli in 10 µl reactions and plated on ampicillin containing LB broth agar plates. After performing colony PCR for size validation with universal primers adjacent to the pUC19 insertion site (M13-for: 5’-GTAAAACGACGGCCAG-3’; M13-rev: 5’-CAGGAAACAGCTATGAC-3’), 3 – 5 ml ampicillin-containing bacterial cultures of preferred clones were incubated at 37 °C overnight. Plasmids were purified using ZymoPURE Plasmid Mini Prep Kit (Zymo Research). Sequence validation of HDR donor template containing plasmids was performed by Sanger Sequencing (LGC Genomics, Berlin). HA-flanked transgenes were amplified from the plasmids by PCR using the KAPA HiFi HotStart 2x Readymix (Roche) with reaction volumes > 500 µl. PCR products were purified and concentrated using paramagnetic beads (AMPure XP, Beckman Coulter). This purification process included two washing steps in 70 % ethanol. Resulting HDRT concentrations were adjusted to 2 µg/µl in nuclease free water. For quantification, HDRTs were diluted 1:20 and measured on Nanodrop 1000 (Thermo Fisher). For optimal quantification accuracy, we recommend using Qubit fluorometer or similar.

### Electroporation buffer

100 ml of electroporation buffer 1M were made according to the original publication^37^ by dissolving 37.3 mg KCl, 142.8 mg MgCl_2_ and 910 mg Mannitol in 40 ml sterile ddH_2_O (Ampuwa) prior to adding 60 ml of 0.2 M Phosphate Buffer Solution (Na_2_HPO_4_/NaH_2_PO_4_; pH 7.2). The buffer was sterile filtered using 0.2 µm syringe filters, aliquoted and stored at - 20°C. Prior to use it was thawed, shaken to dissolve intermittently formed precipitates and placed on ice.

### Formulation of ribonucleoproteins and mix with dsDNA templates

Per transfection of 10^6^ primary human T cells, 0.5 µl of an aqueous solution of 15–50 kDa poly(L-glutamic acid) (PGA, Sigma-Aldrich, 100 µg/µl) was mixed with 0.48 µl of synthetic modified sgRNA (2’-O-Methyl at 3 first and last bases, 3’ phosphorothioate bonds between first 3 and last 2 bases; Synthego, 3.2 µg/µl = 100 µM in TE Buffer; in some experiments modified sgRNA was provided by IDT) by pipetting thoroughly. 0.4 µl Recombinant *Streptococcus pyogenes* Cas9 protein (Alt-R S.p. Cas9 Nuclease V3, IDT, 10 µg/µl = 61 µM) was added and again mixed by thorough pipetting. The molar ratio of Cas9 and sgRNA was thus 1:2. The mixture was incubated for 15 minutes at room temperature (RT) to allow for RNP formation and placed on ice afterwards. For DNA-escalation studies, 0.25-2.0 µl of HDRT (stock concentration: 2 µg/µl) per 10^6^ cells was added just prior to electroporation. For all other experiments, 0.5 µl HDRT per 10^6^ cells was used.

### Electroporation

Anti-CD3 and anti-CD28 stimulated primary human T cells were resuspended, pooled and washed twice in sterile PBS at 100 g for 10 minutes, RT. Afterwards, they were resuspended in 20 µl / 10^6^ cells ice-cold electroporation buffer (1M, or P3 (Lonza) as indicated). The exposure time to the electroporation buffers was kept as short as possible. For electroporation of 1× 10^6^ cells, 20 µl of resuspended cells were transferred to 1.88 µl of RNP/HDRT (except during DNA-escalation studies where different volumes were used) and mixed thoroughly. Afterwards, the T cell / RNP / HDRT mixture was transferred onto a 16-well electroporation strip (20 µl = 10^6^ cells per well, Lonza) or an electroporation cartridge (100 µl = 5×10^6^ cells, Lonza). Cells were carefully transferred onto the electroporation strips using 200 µl tips to avoid trapping air in the solution. Electroporation strips and cartridges were tapped onto the bench several times to ensure correct placement of fluids within the electroporation vessel. Electroporation was performed on a 4D-Nucleofector™ Device (Lonza) using the program EH-115. Directly after electroporation, pre-warmed T cell medium was added onto the cells (90 µl per well and 450 µl per cartridge). Afterwards, resuspended cells were transferred to 96-well round bottom plates (50 µl / well) containing 150 µl pre-warmed T cell medium per well (with or without HDR-enhancing supplements) at a density of 0.5× 10^6^ cells per well. For large scale CAR-T cell generation by electroporation in cartridges, 950 µl of pre-warmed T cell medium was used for initial resuspension, and cells were transferred into 24-well flat bottom plates (500 µl / well) containing 1.5 ml of T cell medium (with or without HDR-enhancing supplements) at a density of 2.5 × 10^6^ cells per well.

### Lentivirus production and transduction

Propagation of lentiviruses for CD19.2-CAR overexpression were prepared as previously described using transient transfection of 293T cells^57^. Briefly, the CD19.2 construct was cloned into the SIN epHIV7 lentiviral vector plasmid. Supernatant from transfected 293T cells was filtered and concentrated by ultra-centrifugation. Titers were determined by transduction of H9 cells with a dilution series of concentrated lentivirus from supernatants. For T cell transduction, primary human T cells were enriched and plated on anti-CD3/CD28-coated tissue culture plates in T cell medium. After two days, transduction was performed by centrifugation at 800 g for 30 minutes at 32°C with lentiviral supernatant (MOI of 1) supplemented with 1 mg/mL protamine sulfate (APP Pharmaceuticals, Barceloneta, Puerto Rico). Three days later, transduced T cells were sub-cultured in G-Re×10 devices (Wilson Wolf).

### CAR T cell expansion

First medium change or first splitting of cells was performed 18 hours after electroporation unless stated otherwise. Cells were expanded in T cell medium on 96-well round bottom plates. During large scale CAR T cell generation for subsequent functional characterization, G-Re×10 devices (Wilson Wolf) were used for expansion. On day 14 after blood collection, cell products were frozen in FCS supplemented with 10% DMSO at a density of 10^7^ cells / ml.

### Drug treatments before and after nucleofection

To test the individual effects of selected DNA-sensor inhibitors, inhibitors were added to the T cells cultured in 24-well stimulation plates at the following concentrations 6-8 hours prior to electroporation: ODN A151 (Invivogen): 5 µM; RU.521 (Invivogen): 9.64 µM; H151 (Invivogen): 3.58 µM; BX795 (Invivogen): 5 µM. The medium was mixed by pipetting and the resuspended T cells were incubated at 37°C and 5% CO_2_ until electroporation.

Combined DNA-sensor inhibition: six hours prior to electroporation, 1 ml of the T cell medium per well (50 % of the medium) was removed from the 24-well stimulation plates and 10 µl of the ODN A151 and 1 ul RU.521 was added to the remaining medium in the stimulation plate at the following concentrations: ODN A151: 5 µM; Ru.521: 4.82 µM. The medium was mixed by pipetting and the resuspended T cells were incubated at 37 °C and 5 % CO_2_ until electroporation.

To test the effects of HDR-enhancing substances (Alt-R HDR Enhancer (HDR-Enhancer version 1 [V1]), and HDR Enhancer version 2 [V2], both available at Integrated DNA Technologies) electroporated and resuspended T cells were transferred onto cell culture plates containing pre-warmed T cell medium with HDR-enhancing supplements in a volume ratio of 1:3 (50 µl cell suspension were added to 150 µl supplemented T cell medium). Concentrations tested ranged from 3.25 µM to 30 µM for HDR-Enhancer v1, and from 0.25 µM to 2 µM for HDR-Enhancer v2. All concentrations stated in this publication refer to the supplemented T cell medium before seeding of T cells.

### Multiplex cytokine analysis of cell culture supernatants

To characterize the cytokine response to transfected DNA, 100 µl of supernatant were harvested 24 hours after transfection. Supernatants were transferred to a sterile 96-Well-Plate, sealed and stored at −80 °C until analysis. After thawing of supernatants, measurements were performed using the human Proinflammatory Panel 1 kit (for cytokines such as IL-1β, IL-6, IL-10, IL-13, TNFα) from Meso Scale Diagnostics according the manufacturer’s instructions. Analysis was performed at the Immunological Study Lab of CheckImmune GmbH on a Mesoscale Discovery platform. For each sample, the respective value of the electrochemiluminescence (RFU-Signal) of the analyte concentration was calculated on the basis of a calibration curve and the blank control values were subtracted (fresh T cell medium). Measurements of the Meso Scale Diagnostics assay were performed and evaluated in accordance with the International Conference on Harmonisation Good Clinical Practice Guideline under ‘Validation of analytical procedures’.

### Flow cytometry

Unless stated otherwise, flow cytometric assessments were carried out on a Cytoflex LX device (Beckman Coulter) using the 96-well plate format. Measurements of cell concentrations / cell counts were performed in 96-well flat-bottom cell culture plates, other measurements were performed in 96-well round-bottom cell culture plates. All staining panels are specified in **Suppl. Table 2**. Representative gating strategies for flow cytometry panels are depicted in **Suppl. Fig. 7**. Cell concentrations were assessed by acquiring 20 µl of resuspended cells diluted 1/10 in PBS without any prior washing steps.

For detection of CAR-expression after knock-in, 40 µl of cell suspension was transferred onto the 96-U-Bottom-well plate and assessed following a series of successive washing and staining steps. For phenotyping, approximately 100,000 T cells were aliquoted per well. Each washing step included adding 240 µl PBS, centrifuging the plates at 400 g, 5 minutes, RT, discarding the supernatants and resuspending the pellets in the remaining volume by vortexing briefly. For any individual staining procedure, a mastermix of the antibodies / dyes diluted in PBS was prepared. 20 µl of the mastermix were added per well. The plates were vortexed briefly, incubated for 15 minutes at 4°C and briefly vortexed again. Because anti-Fc (anti-IgG1, Fc_γ_ part) antibodies used to detect CAR protein can potentially bind to other detection antibodies, a first extracellular staining step with anti-Fc antibody only and a Live-Dead discriminating dye, and subsequent washing step were performed prior to any other staining steps. The phenotype assessment presented in **Suppl. Fig. 6 a, b** was performed on a LSR-II Fortessa flow cytometer (BD Biosciences). The staining for this experiment was performed in 5 ml FACS tubes (Corning). Apart from this difference, it was carried out analogously to the other staining procedures.

### Cell cycle analysis

Stimulation of CD3-enriched T cells as well as DNA-sensor inhibition treatment with RU.521 and ODN A151 were performed as described above. For the staining, 200,000 cells were harvested, washed and stained with Live/Dead Fixable Blue (Invitrogen) for discrimination of dead cells. Then, cells were stained as described previously^39^. Briefly, the cells were fixed and permeabilized using the Intracellular Fixation & Permeabilization Buffer Set (Thermo Fisher, 88-8824). A mastermix containing 7AAD and KI67 Alexa Fluor 647 (Biolegend) was added to the cells for 30 minutes at 4 °C in the dark. After washing, cells were resuspended in PBS and run on the Cytoflex LX cytometer (Beckman Coulter). The 7AAD-dye stains DNA and indicates individual cell genomic DNA content to allow for differentiation of cells in resting (G0) and G1-phase vs. cells in S- and G2-phase. Ki-67 is an activation and proliferation marker which indicates cells outside resting phase (G0).

### VITAL assay to assess cytotoxicity

Effector T cells *(TRAC*-CD19.1-CAR T cells; *TRAC*-BCMA-CAR T cells and unedited (wildtype) T cells; each from the same 3 donors) were co-cultured with target cells expressing the surface protein recognized by the CAR, and control cells negative for this marker. For the CD19 VITAL assay, Nalm6 cells (CD19+, BCMA-) engineered to express Green Fluorescent Protein (GFP) served as target cells and CD19-Knock-Out Nalm6 cells engineered to express Red Fluorescent Protein (RFP) served as control cells. For the BCMA VITAL assay, MM.1S (CD19-BCMA+) engineered to express GFP served as target cells, while the same control cells were used. A 1:1 suspension of target and control cells was added to 25,000 (CAR) T cells in 96-well round bottom cell culture plates at 3 different effector : target : control cell ratios (1 : 2 : 2, 1 : 1 : 1 and 1 : 0.2 : 0.2). The plates were centrifuged at 100 g, 3 min, RT, and then incubated at 37 °C, 5 % CO_2_ for 4 hours. Afterwards, the plates were centrifuged at 400 g, 10 min, RT, and the supernatant was discarded. Cell pellets were resuspended in PBS and the plates centrifuged again at 400 g, 10 min, RT. The supernatant was discarded and the cells were stained with LIVE/DEAD^TM^ Fixable Blue Dead Cell Stain (Invitrogen, L23105) diluted 1:60 in 20 µl PBS per well for 10 min at 4°C. Afterwards, 240 µl PBS / well were added and the plates were centrifuged at 400 g, 15 minutes, RT. The supernatant was discarded, the cells were resuspended in 50 µl PBS and analysed by flow cytometry. Effector cell mediated cytotoxicity was calculated from shifts in the target:control cell ratio relative to control conditions without effector cells. The experiment was performed on day 12 after blood collection (day 10 after electroporation).

### Intracellular cytokine analysis

CAR T cells cryopreserved on day 14 after peripheral blood collection (day 12 after electroporation) were thawed by washing twice in pre-warmed RPMI1640 medium supplemented with 20% FCS and 10 ng/mL DNAse I (Roche, 11284932001). Thawed CAR T cells were rested in cytokine-free complete medium (RPMI1640 medium containing 10% FCS and 1% penicillin-streptomycin) overnight at 37 °C, 5 % CO_2_. Nalm6 and Jurkat cells were labelled with 2.5 μM Carboxyfluorescein diacetate succinimidyl ester (CFDA-SE, ThermoFisher Scientific, V12883) and rested in complete medium overnight. 100,000 CAR T cells were co-cultured with CFSE-labelled Nalm6 or Jurkat cells (ratio 1:1) in 96-well round bottom plates for 6 hours. CAR T cells stimulated with 10 ng/mL PMA (Sigma Aldrich, P8139-1MG) and 2.5 μg/mL Ionomycin (Sigma Aldrich, I3909) were included as positive control for polyclonal stimulation independent of TCR expression, and CAR T cells cultured alone were included as negative control. Brefeldin A (Sigma Aldrich, B5936) was added at a concentration of 10 μg/mL after 1 hour of co-culture/stimulation. After 6 hours of stimulation, cells were harvested and stained with fixable blue dead cell stain dye. Subsequently, cells were fixed and permeabilized using the Intracellular Fixation & Permeabilization Buffer Set (Thermo Fisher, 88-8824). A first intracellular staining with anti-Fc AF647 was performed prior to washing twice with 1X permeabilization buffer. A second intracellular staining was performed with all other antibodies as stated in **Suppl. Table 2**.

### Data analysis, statistics and presentation

Flow cytometry data was analysed with FlowJo software v10 (BD). Data from different assays was sorted in Excel (Microsoft). Graphs were created using Prism 9 (GraphPad). Conditions with failed electroporation (indicated by 4D-Nucleofector™ Device) were recorded during experiment and later excluded from analysis. Exploratory statistics were performed with Prism 9 (Graphpad). Schemes and graphs in the presented Figures were created using an academic license of BioRender (www.biorender.com).

