## Supplementary Figures for "Fast, efficient and virus-free generation of *TRAC*-replaced CAR T cells"

### **Suppl. Fig. 1 - GUIDE-seq indicates reduced off-targets by target-sequence modified *TRAC* sgRNA**

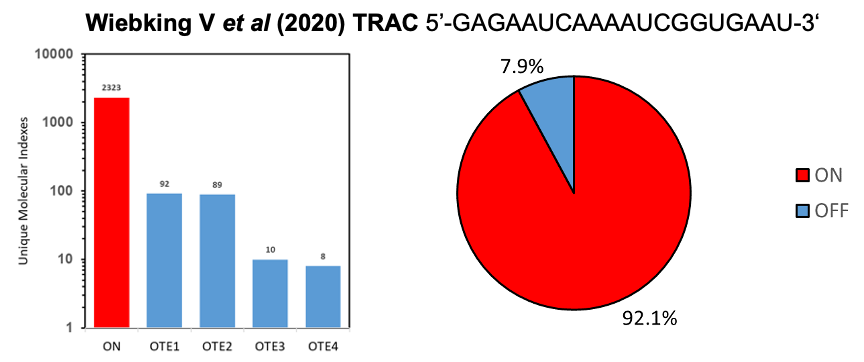

| **TRAC sgRNA (Wiebking 2020)** | **Reads** | **Target Sequence** | **Matched sequence (with highlighted mismatches to sgRNA)** | **Aligned sequence** | **Leveshtein distance** | **Chromo-**  **some** | **Alignment start** | **Alignment stop** | **strand** |
| --- | --- | --- | --- | --- | --- | --- | --- | --- | --- |
| On-target | 2323 | GAGAATCAAAATCGGTGAAT | GAGAATCAAAATCGGTGAAT | ATTCACCGATTTTGATTCTC | 0 | chr14 | 22547575 | 22547595 | - |
| Off-target event 1 | 92 | GAGAATCAAAATCGGTGAAT | TCTTATCAAAATCAGTGAAT | ATTCACTGATTTTGATAAGA | 5 | chr6 | 1112737 | 1112757 | - |
| Off-target event 2 | 89 | GAGAATCAAAATCGGTGAAT | GTGTATTTAAATTGTTTCAT | ATGAAACAATTTAAATACAC | 8 | chr1 | 237327485 | 237327505 | - |
| Off-target event 3 | 10 | GAGAATCAAAATCGGTGAAT | AAAAAACTAAATCAATTTAT | ATAAATTGATTTAGTTTTTT | 8 | chr22 | 34169261 | 34169281 | - |
| Off-target event 4 | 8 | GAGAATCAAAATCGGTGAAT | TATAATGAAAATTTAAAAAT | TATAATGAAAATTTAAAAAT | 8 | chr8 | 115314523 | 115314543 | + |

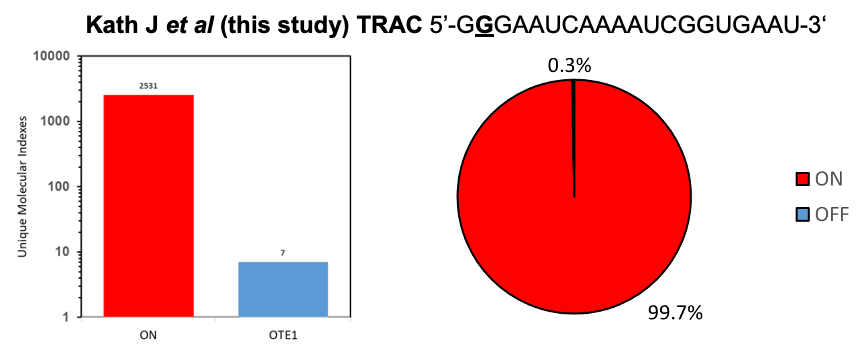

| **TRAC sgRNA (Charité)** | **Reads** | **Target Sequence** | **Matched sequence (with highlighted mismatches to sgRNA)** | **Aligned sequence** | **Leveshtein distance** | **Chromo-**  **some** | **Alignment start** | **Alignment stop** | **strand** |
| --- | --- | --- | --- | --- | --- | --- | --- | --- | --- |
| On-target | 2531 | GGGAATCAAAATCGGTGAAT | GAGAATCAAAATCGGTGAAT | ATTCACCGATTTTGATTCTC | 1 | chr14 | 22547575 | 22547595 | - |
| Off-target event 1 | 7 | GGGAATCAAAATCGGTGAAT | GGAAATATAAATTATTATAT | ATATAATAATTTATATTTCC | 8 | chr19 | 20782355 | 20782375 | - |

**Suppl. Fig. 1:** Results of GUIDE-seq analysis in HEK293 overexpressing SpCas9 for the original *TRAC* sgRNA (Wiebking et al 2020) and this study (Charité) and the respective on- and off-target event locus sequences which were identified.

### **Suppl. Fig. 2 – Impact of cell number, RNP dose and anionic nanoparticle PGA, stimulation conditions and electroporation buffer on CAR integration into *TRAC* locus**

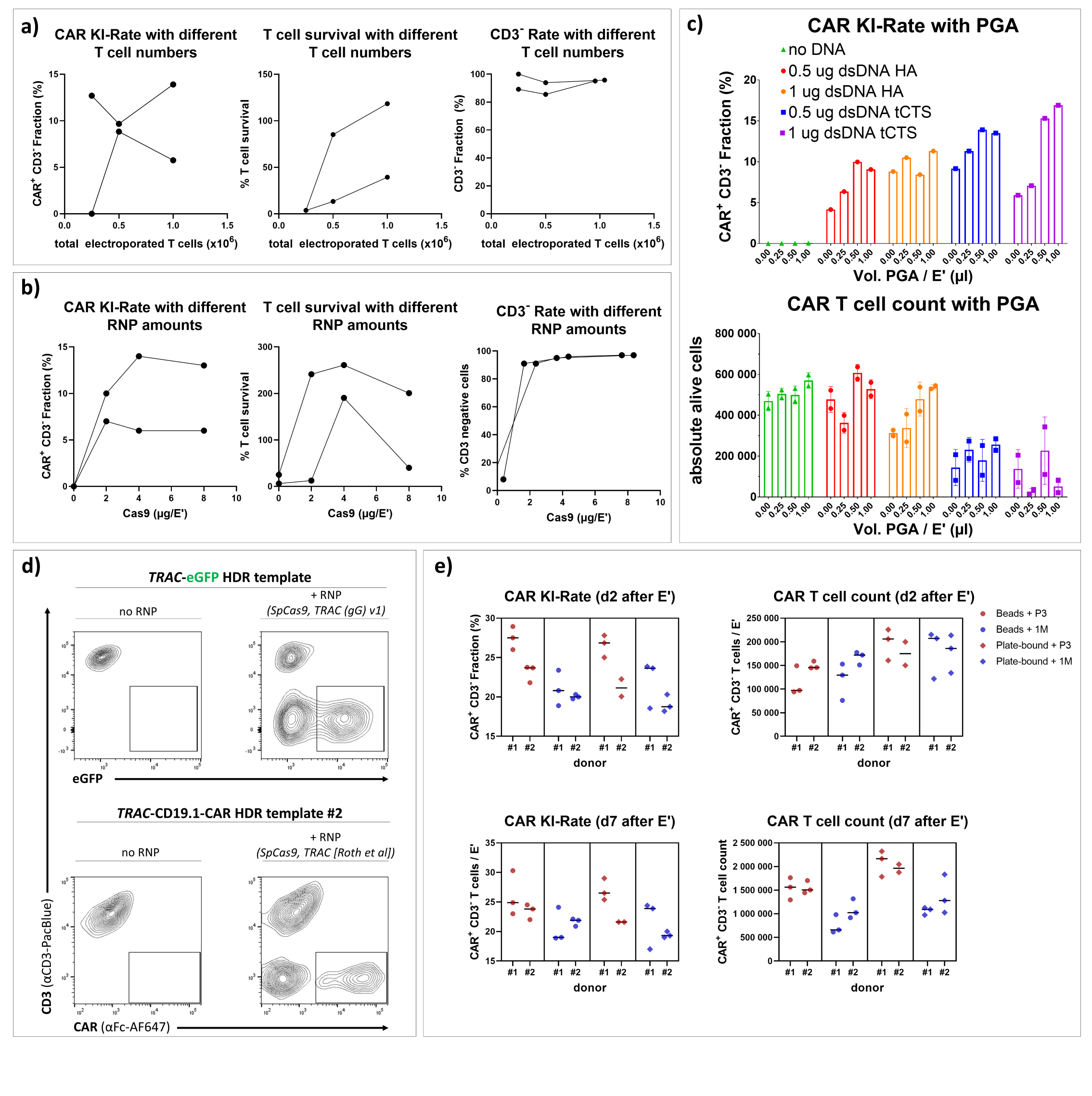

**Suppl. Fig. 2:** Optimization of different aspects of the transfection protocol. (a) Summary of the effect of different cell densities during transfection on CAR integration rate, relative T cell survival to mock-transfected and CD3-knock-out rate as a marker of effective *TRAC* cutting. n = 2 biological replicates. Lines indicate paired results. (b) Summary of the effect of different amounts of RNP during transfection on CAR integration rate, T cell survival relative to mock-electroporated cells, and CD3-knock-out rate as a marker of effective *TRAC* cutting. In contrast to other experiments, RNPs were pre-complexed with sgRNA at 2.5:1 molar ratio to SpCas9 prior to transfection. n = 2 biological replicates. Lines indicate paired results. (c) Summary of the effect of the different amounts of anionic nanoparticle poly-glutamic acid (PGA) on the relative CAR integration rate as well as CAR T cell yield 4 days after transfection. In this experiment, RNPs were formulated by first mixing 0.48 µl sgRNA (3.2 µg/µl) with the respective amount of PGA, prior to adding 0.4 µl of Cas9 (10 µg/µl). n = 1 biological replicate, 2 technical replicates. CAR knock-in rate was only assessed in 1 technical replicate (d) Representative flow cytometry plots of editing results after transfection with dsDNA HDR template alone or together with co-delivered RNP. HDR templates encoding eGFP or a CD19-CAR, and their respective RNP were used (similar setup as in Fig 3 d). (e) Summary of data comparing different stimulation conditions and electroporation buffers and their effect on CAR knock-in rate and CAR T cell yield. Range of y-axes differs for CAR T cell count of d2 after electroporation (E’) and d7 after E’.

### **Supp. Fig. 3 – Cell cycle analysis after combined DNA-sensor inhibition indicates increased proportion of cells within S-phase**

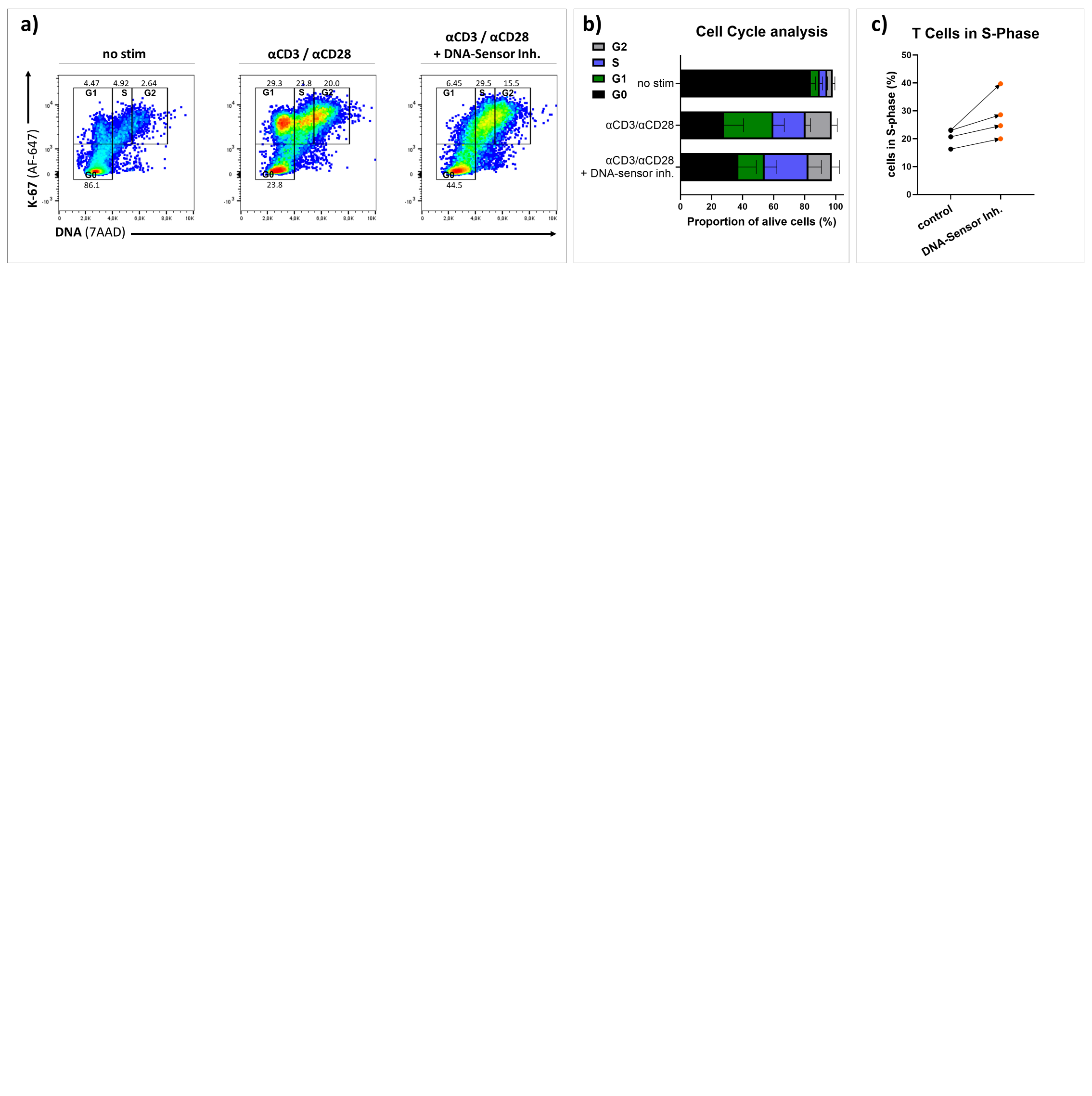

**Suppl. Fig. 3:** Cell cycle analysis after DNA-sensor inhibition. (a) Representative flow cytometry plots of a cell cycle staining performed in 4 biological replicates in two independent experiments. Analysis of unstimulated T cells (left), anti-CD3/CD28 stimulated T cells (middle), and anti-CD3/CD28 T cells after a combined 6-hour treatment with the DNA-sensor inhibitors RU.521 and ODN A151 (right). (b) Summary of a. (c) Summary of a for the impact of DNA-sensor inhibition on cells within S-phase.

**Suppl. Fig. 4 – HDR enhancers improve suboptimal gene editing but are sensitive to temperature and treatment time**

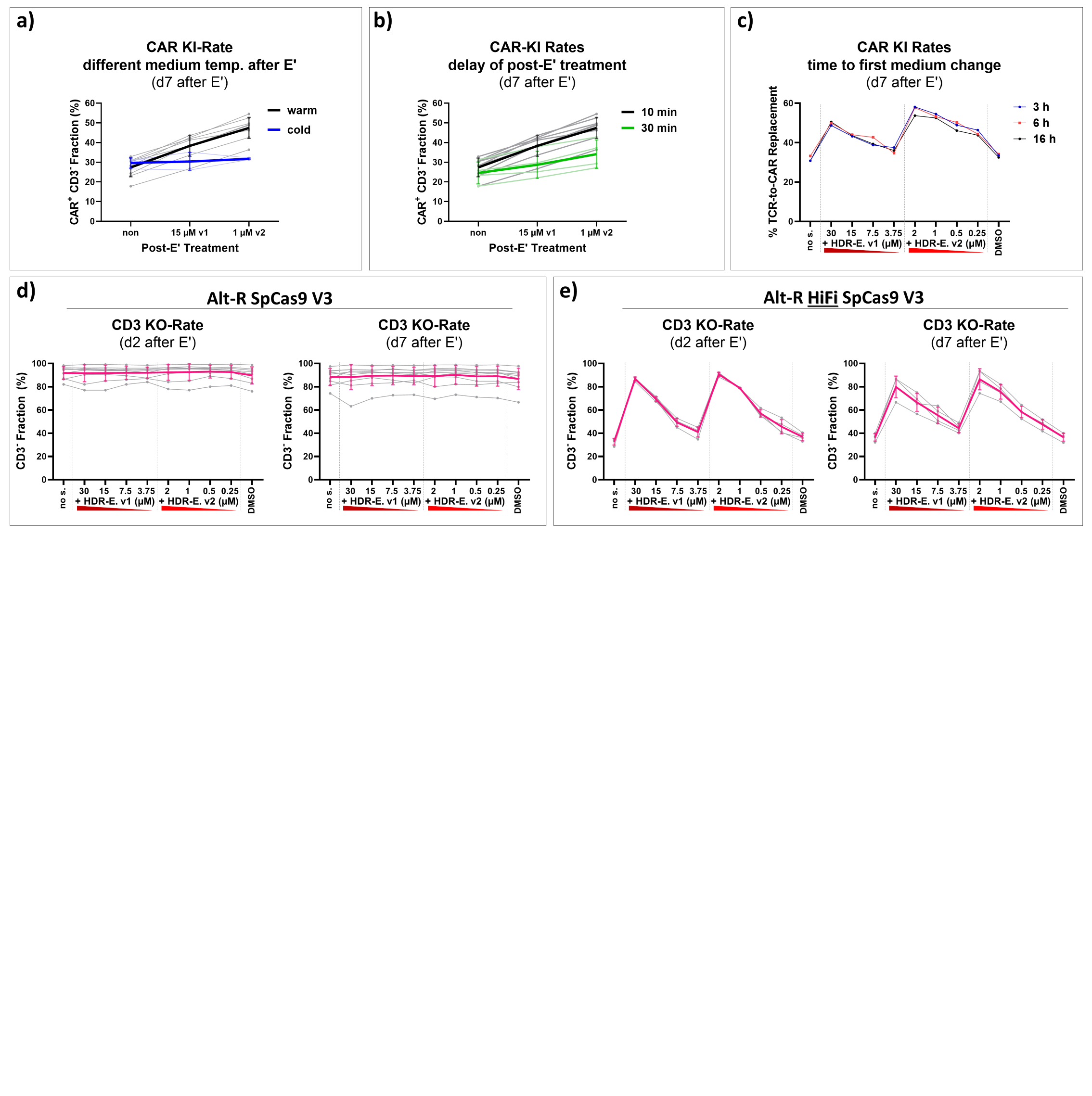

**Suppl. Fig. 4:** Specific aspects of the HDR-enhancers’ mode of action**.** (a) Temperature-dependent effects of HDR enhancers. Black lines represent conditions with T cell medium with/without supplemented HDR enhancers that was pre-warmed to 37°C prior to cell transfer (n = 9 biological replicates). In blue: T cell medium after transfection was supplemented with/without HDR enhancers, but was kept in 4°C until time of transfection and cell transfer (n = 3 biological replicates). (b) Time-sensitive effects of HDR enhancers. Editing outcomes of T cells subjected to a 30 min delay between electroporation (with immediate resuspension in T cell medium) and transfer into HDR-enhancer supplemented medium are shown in green (n = 4 biological replicates). Controls are shown in black (n = 9; same as in a). (c) Effects of HDR-enhancer treatment duration on editing outcomes. At different time points after electroporation and consecutive cell transfer to HDR enhancer supplemented T cell medium, half the supplemented medium was exchanged with drug-free T cell medium. For the 3 h and 6 h conditions, a second medium change was performed 16 h after electroporation. n = 1 healthy donor. (d,e) CD3-knock-out efficiency from experiment presented in Fig. 3 c + f. Data points represent the percentage of CD3^-^ T cells two days after editing using either Alt-R SpCas9 V3 (d) or Alt-R HiFi SpCas9 V3 (e).

### **Suppl. Fig. 5 – Complete data set of experiments comparing different cell treatments prior and/or after transfection as well as increasing amounts of dsDNA donors**

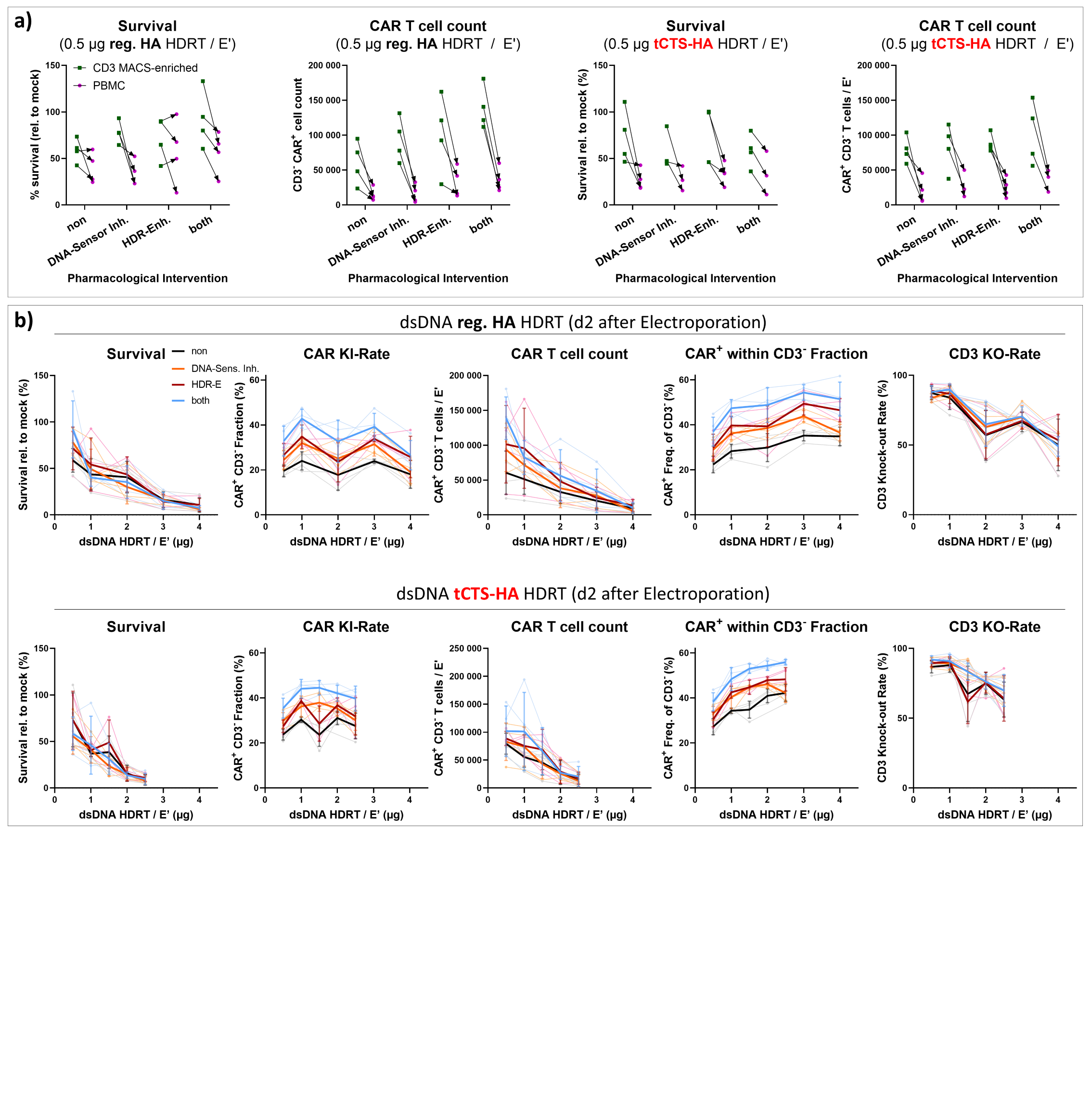

**Suppl. Fig. 5:** (a) Comparison of two different cell populations as starting material for CAR T cell generation. PBMC with or without additional CD3-MACS-enrichment were stimulated on anti-CD3/CD28-coated wells and electroporated with HDRTs (reg. HA or tCTS-HA). Cells were analyzed by flow cytometry two days after electroporation (n = 4 biological replicates in 2 independent experiments). (b) Complete data sets are displayed for the experiment which were partially presented throughout the manuscript, including in Fig. 1 e, Fig. 2 f and Fig. 4 b. These plots comprise the results of all tested parameters for all transfected DNA doses and both HA formats. Cells were analyzed by flow cytometry two days after electroporation (n=4 biological replicates from 2 independent experiments.) These plots do not contain the two additional biological replicates added for reg. HA HDRTs in Fig. 4 b.

### **Suppl. Fig. 6 – TRAC-integration leads to lower CAR expression when compared with other means of transgene overexpression**

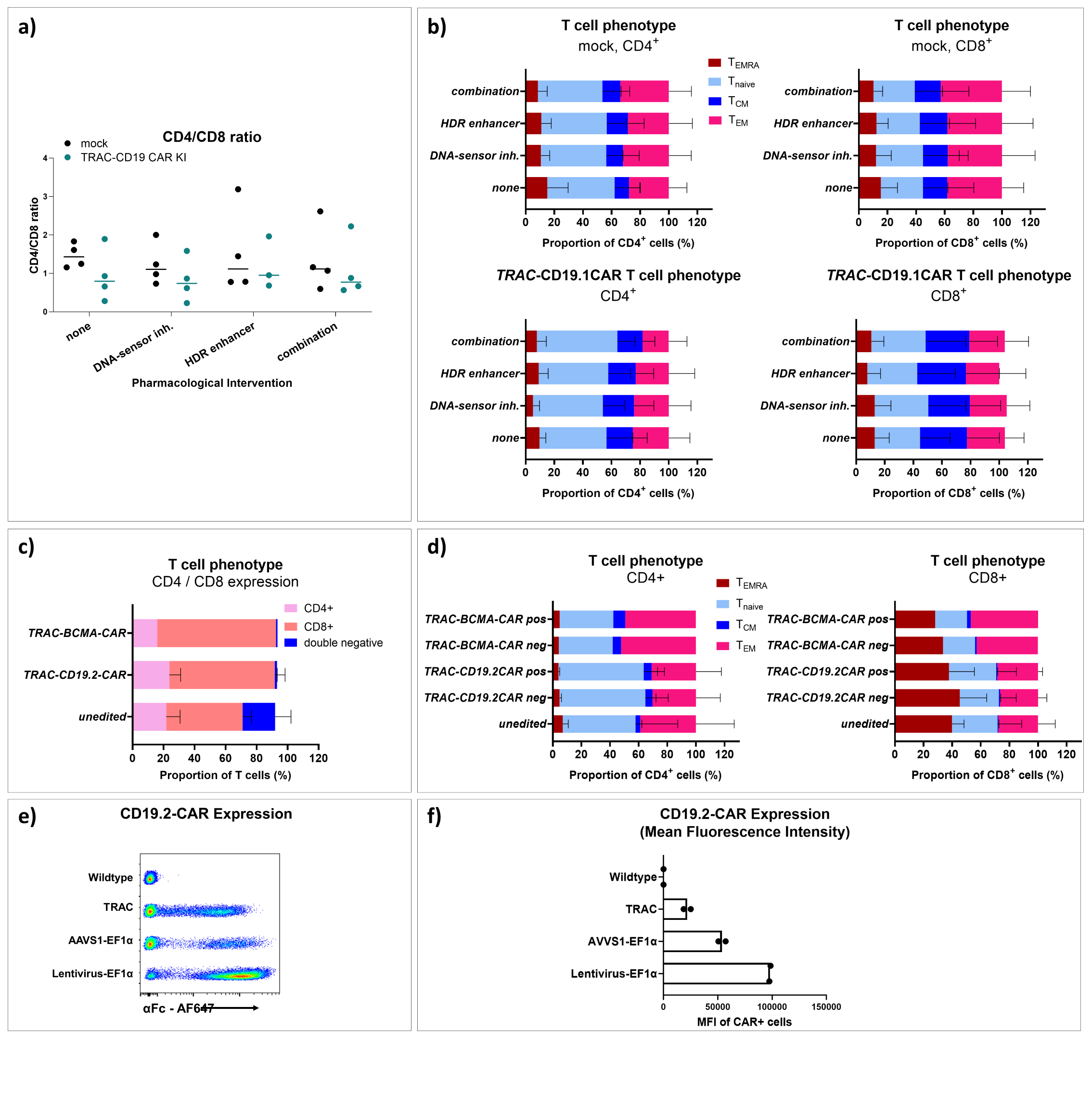

**Suppl. Fig. 6:** (a,b) Cell phenotypes of mock-transfected and *TRAC*-integrated CD19-CAR T cells after receiving different pharmacological interventions. T cells were analyzed by flow cytometry nine days after transfection. (a) CD4:CD8 ratios in different populations (b) Summary of T cell memory phenotypes (T_EMRA_: CD45RA^+^, CCR7^-^; T_naive_: CD45RA^+^ CCR7^+^; T_CM_: CD45RA^-^, CCR7^+^; T_EM_: CD45RA^-^, CCR7^-^) in CD4^+^ CD8^-^ or CD4^-^ CD8^+^ T cells. (c, d) Comparison of T cell phenotypes of CD19.2-CAR T cells (n = 2 biological replicates analyzed in 2 technical replicates) and BCMA-CAR T cells (n=1) 24 hours after thawing. Prior to cryopreservation, T cells were expanded for a total of 14 days. (e, f) Comparison of CD19.2-CAR expression levels of CAR T cells generated using different methods. ‘*TRAC’* indicates the method described in this manuscript. For condition ‘AAVS1-EF1a’, the protocol was adapted to insert a lentivirus-derived overexpression cassette including an EF1α promoter and a poly-adenylation signal into the *AAVS1* safe-harbor locus. Such CAR T generated in a virus-free approach were compared with CD19.2-CAR T cells generated by lentiviral gene transfer.

### **Suppl. Fig. 7 – Representative gating strategies for flow cytometry assays**

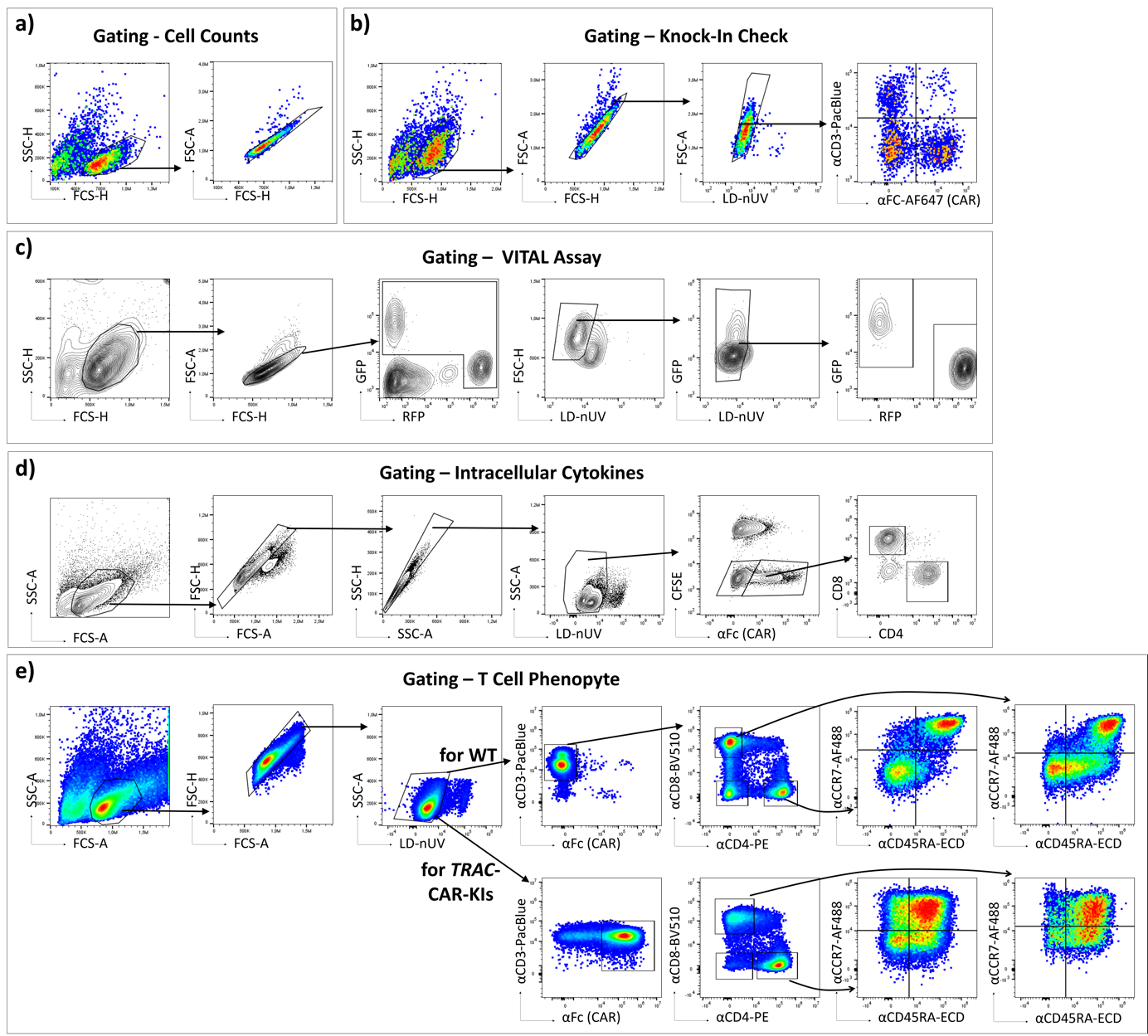

**Suppl. Fig. 7** Representative gating strategies for flow cytometry analyses. (a) Gating for cell count determination. (b) Gating for analysis of editing outcomes (‘Knock-In Check’). Such gating to identify TCR-replaced CAR T cells was applied in all experiments that did not require T cell stimulation or intracellular staining. (c) Gating strategy to determine target:control cell ratios for the VITAL assays (as presented in Fig. 5 b, c). (d) Gating after intracellular staining to determine cytokine production following different modes of (CAR) T cell stimulation (as presented in Fig. 5 e-h). (e) Gating strategy for phenotype assessment of (CAR) T cells (as presented in Suppl. Fig. 6 d)
